# Pan-cancer prediction of tumor immune activation and response to immune checkpoint blockade from tumor transcriptomics and histopathology

**DOI:** 10.1101/2025.06.27.661875

**Authors:** Sumit Mukherjee, Sumeet Patiyal, Lipika R. Pal, Keluo Yao, Tian-Gen Chang, Sumona Biswas, Saugato Rahman Dhruba, Amos Stemmer, Arashdeep Singh, Abbas Yousefi-Rad, Tien-Hua Chen, Binbin Wang, Denis Marino, Wonwoo Shon, Yuan Yuan, Mark Faries, Omid Hamid, Karen Reckamp, Barliz Waissengrin, Beatriz Ornelas, Pen-Yuan Chu, Lisa Ley, Dilara Akbulut, Nourhan El Ahmar, Sabina Signoretti, David A. Braun, Joo Sang Lee, Hyunjeong Joo, Hyungsoo Kim, Arsen Osipov, Robert A. Figlin, Jair Bar, Iris Barshack, Chi-Ping Day, Sridhar Hannenhalli, Karine Sargsyan, Andrea B. Apolo, Kenneth Aldape, Muh-Hwa Yang, Michael B. Atkins, Ze’ev A. Ronai, Danh-Tai Hoang, Eytan Ruppin

## Abstract

Accurately predicting which patients will respond to immune checkpoint blockade (ICB) remains a major challenge. Here, we present TIME_ACT, an unsupervised 66-gene transcriptomic signature of tumor immune activation derived from TCGA (The Cancer Genome Atlas) melanoma data. First, we demonstrate that TIME_ACT scores accurately identify tumors with activated immune microenvironments across different cancer types. Further, analysis of spatial features reveals that tumor microenvironment regions with dense lymphocyte infiltration near tumor cells have high TIME_ACT scores, successfully marking localized immune activation. Second, across 25 transcriptomic ICB cohorts encompassing nine cancer types, TIME_ACT achieves a mean AUC of 0.76 and a mean odds ratio of 5.77, significantly outperforming 30 established transcriptomic signatures and prediction methods for ICB response, including a recently developed foundation model for immunotherapy response prediction. Third, we show that TIME_ACT scores can be accurately inferred from routine tumor histopathology slides and that slide-inferred TIME_ACT scores predict ICB response across nine new independent patient cohorts spanning eight cancer types, achieving a mean AUC of 0.72 and a mean odds ratio of 4.99. These findings establish TIME_ACT as a robust, pan-cancer biomarker that enables accurate, low-cost, and clinically scalable prediction of ICB response from routine histopathology.

## Introduction

Immune checkpoint blockade (ICB) therapies have transformed the landscape of cancer immunotherapy, yielding considerable clinical benefits in a subset of patients across multiple cancer types ^1–3^. Among these, melanoma and non-small cell lung cancer (NSCLC) exemplify a prototypical immunogenic tumor where ICB has demonstrated remarkable therapeutic success ^4,5^. Despite this progress, a substantial fraction of patients do not respond, underscoring an unmet need for predictive biomarkers that can guide treatment decisions and improve patient stratification ^6^. Importantly, a critical unmet challenge is identifying which tumors possess sufficient immune activation to benefit from ICB therapy.

A key determinant of ICB efficacy is the tumor microenvironment (TME), which plays a central role in modulating anti-tumor immunity ^7,8^. TME exhibits considerable heterogeneity and has been broadly categorized as “Hot” and “Cold” immunophenotypes ^9^. Hot tumors are enriched with tumor-infiltrating lymphocytes (TILs), exhibit high expression of immune checkpoints, and demonstrate active inflammatory signaling, robust antigen presentation, and elevated cytotoxic T lymphocyte (CTL) activity, characteristics generally associated with better responses to ICB ^7^. In contrast, cold tumors are immunologically inactive, characterized by sparse immune infiltration, impaired antigen processing, and low expression of interferon-stimulated genes and chemokines, contributing to immune evasion and therapy resistance ^5^. While it is challenging to distinguish between the presence of immune cells and functional immune activity, several studies have attempted to classify hot and cold tumors ^10–16^. Notably, many of these studies have generated gene expression signatures that capture only a single facet of tumor “hotness,” such as inflammation, cytotoxic T-cell infiltration, or interferon signaling, and have often been developed within a single cancer type without testing reproducibility across tumor types and cohorts, thereby limiting their utility as broadly applicable predictive biomarkers. A critical unmet challenge, hence, is the development of a more general biomarker that integrates multiple axes of tumor “hotness” and identifies which individual tumors (independent of tumor type) possess sufficient immune activation to benefit from ICB therapy.

A variety of transcriptomic approaches based on immune-related signatures have been developed to predict ICB response, including IFN-γ-related signatures ^17^, T cell signatures ^18,19^, B cell signatures ^20,21^, and antigen presentation machinery-based scores ^22,23^. Furthermore, several transcriptomics-based methods such as TIDE ^24^, IMPRES ^25^, SELECT ^26^, IPRES ^27,28^, IPS ^29^, COX_IS ^30^, PredictIO ^31^, etc., have been developed to capture tumor-immune dynamics or predict ICB response probabilities. However, these methods often exhibit cohort-specific variable performance, and some rely on assumed mechanistic interpretations of response, without directly measuring the activation state of the TME ^32,33^. Recently, COMPASS ^34^, a pan-cancer foundation model for immunotherapy response prediction, used self-supervised contrastive learning on TCGA transcriptomes to encode each tumor as a 44-dimensional representation of predefined immune concepts, then trained a lightweight classifier on this representation to predict ICB response. Although it generalizes across cancer types and checkpoint therapies, it requires labeled ICB cohorts for adaptation, underscoring the need for more comprehensive validation in independent cohorts to further establish its robustness and clinical generalizability. Thus, there remains an important unmet clinical need for simple, biologically grounded biomarkers that directly quantify tumor immune activation and can be translated to clinically accessible modalities, such as routine histopathology, thereby helping democratize precision oncology.

To address this gap, here we introduce TIME_ACT (Tumor Immune MicroEnvironment ACTivation), a robust and interpretable transcriptomic signature derived from TCGA melanoma samples using converging evidence from three previously established and complementary measures of immune activation, encompassing immune infiltration, inflammatory signaling, and histopathology-derived spatial patterns of tumor-infiltrating lymphocytes (TILs). Inferred in melanoma, where ICB treatments have been the most successful to date, TIME_ACT is designed to reflect functional immune engagement rather than the mere presence of immune cells. We first demonstrate that TIME_ACT generalizes to several cancer types beyond melanoma, enabling pan-cancer identification of immunologically hot tumors. We then show that TIME_ACT robustly predicts ICB response across 25 cohorts spanning nine cancer types, outperforming nine established transcriptomics-based prediction methods, 21 published ICB response signatures, and the recently developed pan-cancer foundation model. We then leveraged Path2Omics ^35,36^, our recently developed deep learning model designed to predict transcriptomics from tumor H&E slides, to infer TIME_ACT scores directly from patients’ histopathology slides. We show that the slide-inferred TIME_ACT score successfully predicts patient response to ICB therapies across nine multi-centric independent unseen datasets across eight cancer types. These results underscore the clinical translational potential of TIME_ACT to advance precision oncology in a timely, widely accessible manner using standard pathology slides, offering an exciting way to facilitate patient matching to ICB therapy going forward.

## Results

### Study design and datasets

We designed a multi-step computational framework to identify a transcriptomic signature of tumor immune activation. We then evaluated its predictive utility across both transcriptomic and histopathology data modalities. Firstly, we stratified TCGA melanoma tumors into immune “Hot” and “Cold” groups based on consensus scoring across immune infiltration, inflammation, and TIL patch density, followed by independent pathologist verification, forming the basis for downstream gene signature derivation **(Fig. 1a)**. In the second step, we identified genes upregulated in Hot tumors and enriched for immune-related pathways, resulting in a 66-gene expression signature termed *TIME_ACT*. This signature was then applied to additional TCGA tumor types to assess pan-cancer relevance. Then, its ICB response prediction ability was evaluated in 25 independent transcriptomic cohorts of ICB-treated patients, spanning nine different cancer types, with a total of 1,196 patient samples. Further, to characterize the biological basis of TIME_ACT, we performed co-expression network analysis, single-cell RNA-seq profiling, and spatial transcriptomics **(Fig. 1b)**. In the third step, we extended this framework to histopathology using Path2Omics ^36^, an advanced deep learning approach that builds on our previously established DeepPT framework ^35^. Path2Omics models can infer transcriptome-wide gene expression directly from H&E-stained whole-slide images. We applied the pre-trained Path2Omics models to infer the gene expression in tumor slides from nine new multi-centric unpublished ICB patients’ cohorts spanning eight cancer types with a total of 456 patient slides, TIME_ACT scores were subsequently computed from the slide-inferred transcriptomes to predict response **(Fig. 1c)**. To our knowledge, this study represents the first computational framework predicting ICB response from routine pathology slides across multiple cancer types in an unsupervised approach. Notably, Path2Omics was trained on the TCGA cohort and applied as-is to the treatment cohorts without any additional fine-tuning, thereby mitigating the risk of overfitting.

**Fig. 1.**
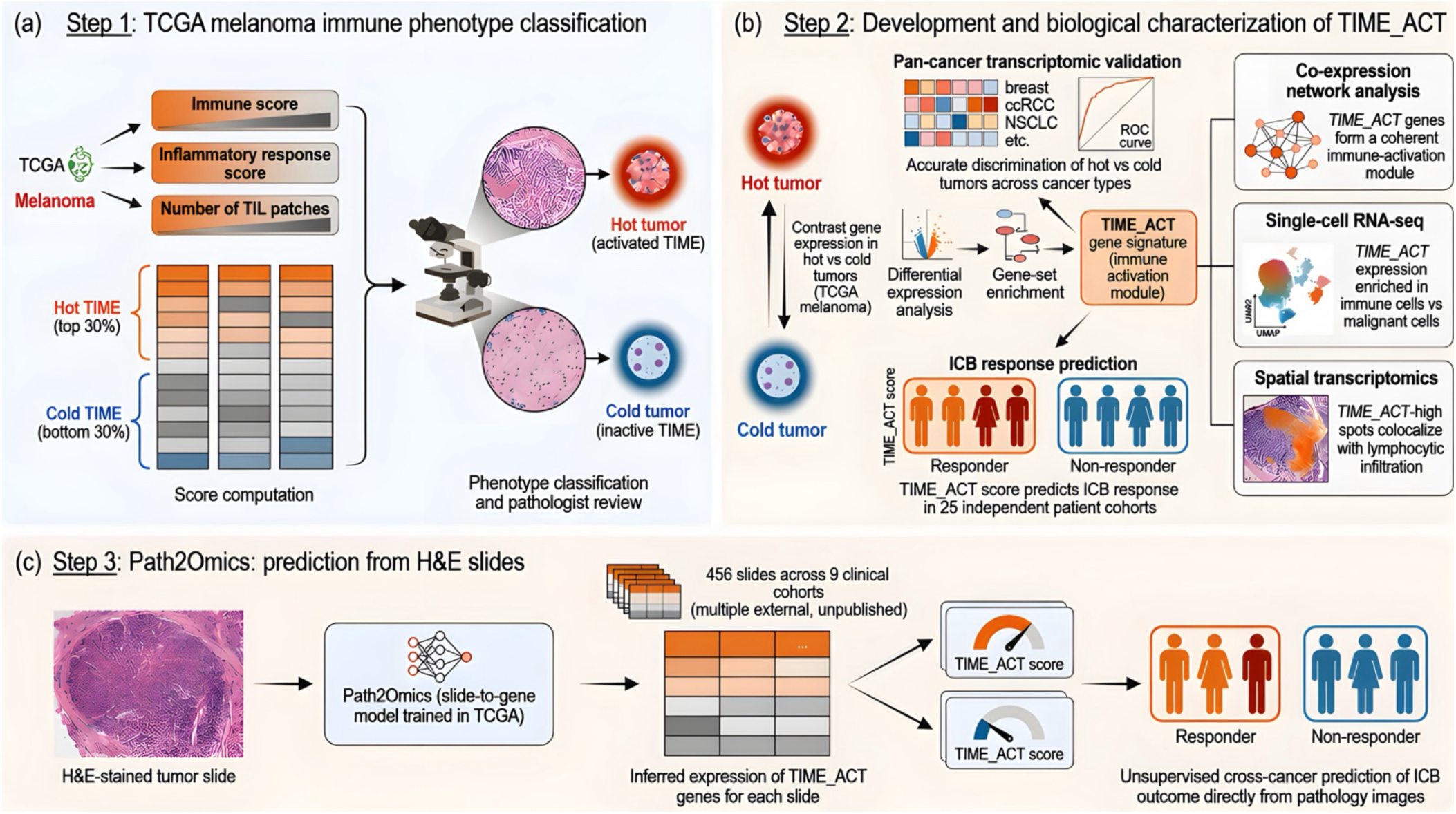
Multi-modal analytical framework for developing and validating the TIME_ACT immune activation signature and predicting ICB response. (a) Schematic overview of immune phenotype classification in TCGA melanoma using three orthogonal immune features: Immune Score, Inflammatory Response Score, and Number of TIL patches, followed by pathologist verification. (b) TIME_ACT’s ability to identify ‘hot’ vs ‘cold’ labeled tumors was identified across additional TCGA cancer types on which it was never trained. Its ability to predict ICB response was then evaluated in 25 independent transcriptomic cohorts of ICB-treated patients. To further investigate the biological basis of TIME_ACT, we performed co-expression network analysis, single-cell RNA-seq profiling, and spatial transcriptomics to uncover its cell-type specificity and spatial localization within the tumor microenvironment. (c) To extend TIME_ACT ICB prediction to histopathology, we applied Path2Omics, a deep learning framework trained on TCGA slides, to infer gene expression directly from H&E-stained images. TIME_ACT scores computed from the slide-inferred expression of TIME_ACT genes were then used to predict ICB response across nine unpublished, unseen clinical cohorts, enabling the first-of-its-kind unsupervised cross-cancer prediction of ICB outcome directly from pathology images.

### Identifying the TIME_ACT signature via an integrative analysis in the TCGA melanoma cohort

To identify tumor immune activation signatures, we first selected melanoma as our discovery cohort due to its well-characterized immunogenicity and relatively high response rate to ICB therapies ^12,40^. To generate a multi-dimensional composite representation of immune activation, we integrated three existing orthogonal metrics of immune activity in TIME that collectively represent immune cell abundance, functional activation, and spatial distribution. Specifically, we used (i) the Immune Score ^37^, to quantify the extent of immune infiltration; (ii) the Inflammation Score ^38^, to measure cytokine-driven immune activation; and (iii) the number of TIL Patches ^39^, to assess the spatial organization of immune-enriched regions within the tumor. Together, these three metrics capture overall levels and spatial dimensions of immune engagement (**Supplementary Fig. 1a**). We found that 363 TCGA-SKCM (melanoma) samples had complete data for all three scores. Then, we used an aggregate ranking across all three metrics for each tumor in the TCGA melanoma cohort (see Methods for details) to assess their overall immune activation. Correlation analysis revealed that immune and inflammation Scores were strongly correlated (Spearman r = 0.8, *p = 1.86e-82*), consistent with the biological coupling between immune cell infiltration and cytokine-driven activation. In difference, their correlations with the number of TIL Patches were moderate (Immune Score: r = 0.40, *p = 9.48e-16*; Inflammation Score: r = 0.41, *p = 3.31e-16*), (**Supplementary Fig. 1b**). These results suggest the while correlated, combining Immune and Inflammation Scores together may provide a more comprehensive measure of overall immune infiltration, and the inclusion of TIL Patches adds a complementary important spatial context for estimating the overall immune activation. Tumors in the top 30% across these metrics were defined as *immune-active (“Hot”)*, and those in the bottom 30% as *immune-inactive (“Cold”)* **(Fig. 1a)**. This approach identified 66 Hot and 49 Cold tumors, yielding a total of 115 tumors, representing 32% of the samples in the cohort, providing an adequate sample size while maintaining stringent classification to enable robust downstream comparisons between tumors with clearly active or inactive immune landscapes. The middle 40% of the TCGA samples were excluded to avoid phenotype ambiguity and to ensure a stringent contrast between Hot and Cold groups. We note that the resulting signature is used in all following analyses to quantify the hotness level of any input tumor sample.

Further, to validate the immune Hot and Cold tumor assignments derived from consensus immune scoring, we performed an independent pathology-based assessment using TCGA-SKCM H&E whole-slide images. Tumors classified as immune “Hot” showed high concordance with histologic evidence of lymphocytic infiltration, with 52 of 66 cases confirmed as immune-active. In contrast, 45 of 49 tumors classified as immune “Cold” were confirmed as immune-inactive based on TIL patterns. Consistent with this assessment, the image-derived TIL Patch metric from Saltz et al. ^39^ accurately distinguished immune Hot tumors, achieving an AUC of 0.90 **(Supplementary Fig. 1c)**. These results establish the Hot and Cold tumor groups in TCGA-SKCM as pathologically supported immune phenotypes, and the pathologist-validated subset of 52 Hot and 45 Cold tumors was used as ground truth for subsequent downstream analyses for the derivation of tumor immune-activation signature.

Differential gene expression (DGE) analysis comparing Hot and Cold tumors identified 339 genes that were significantly highly upregulated in Hot tumors **(Supplementary Table 1)**, filtered using stringent thresholds (the highest quartile of upregulated genes in DGE analysis selected based on both log₂FC and Bayesian B-value: log₂FC > 3.344 and B > 36.224) **(Fig. 2a)**. Furthermore, functional enrichment analysis revealed that these genes were significantly involved in immune-related biological processes such as T cell activation, lymphocyte differentiation, etc. **(Supplementary Table 2)**. We further refined this gene list by selecting only those involved in the top 10 most significantly enriched immune-related processes (adjusted p-value < 0.05) **(Fig. 2b)**. This filtering yielded a curated set of 66 genes, which we defined as the “*TIME_ACT”* signature **(Supplementary Table 3)**. Further, analyzing two single-cell RNA-seq (scRNA-seq) datasets from melanoma patient samples ^40,41^ demonstrated that TIME_ACT genes are predominantly expressed in immune cell populations, particularly T cells and B cells, while showing minimal expression in malignant cells, fibroblasts, and cancer-associated fibroblasts (CAFs) **(Fig. 2c)**. This immune cell specificity reinforces the signature’s relevance to the TIME.

**Fig. 2.**
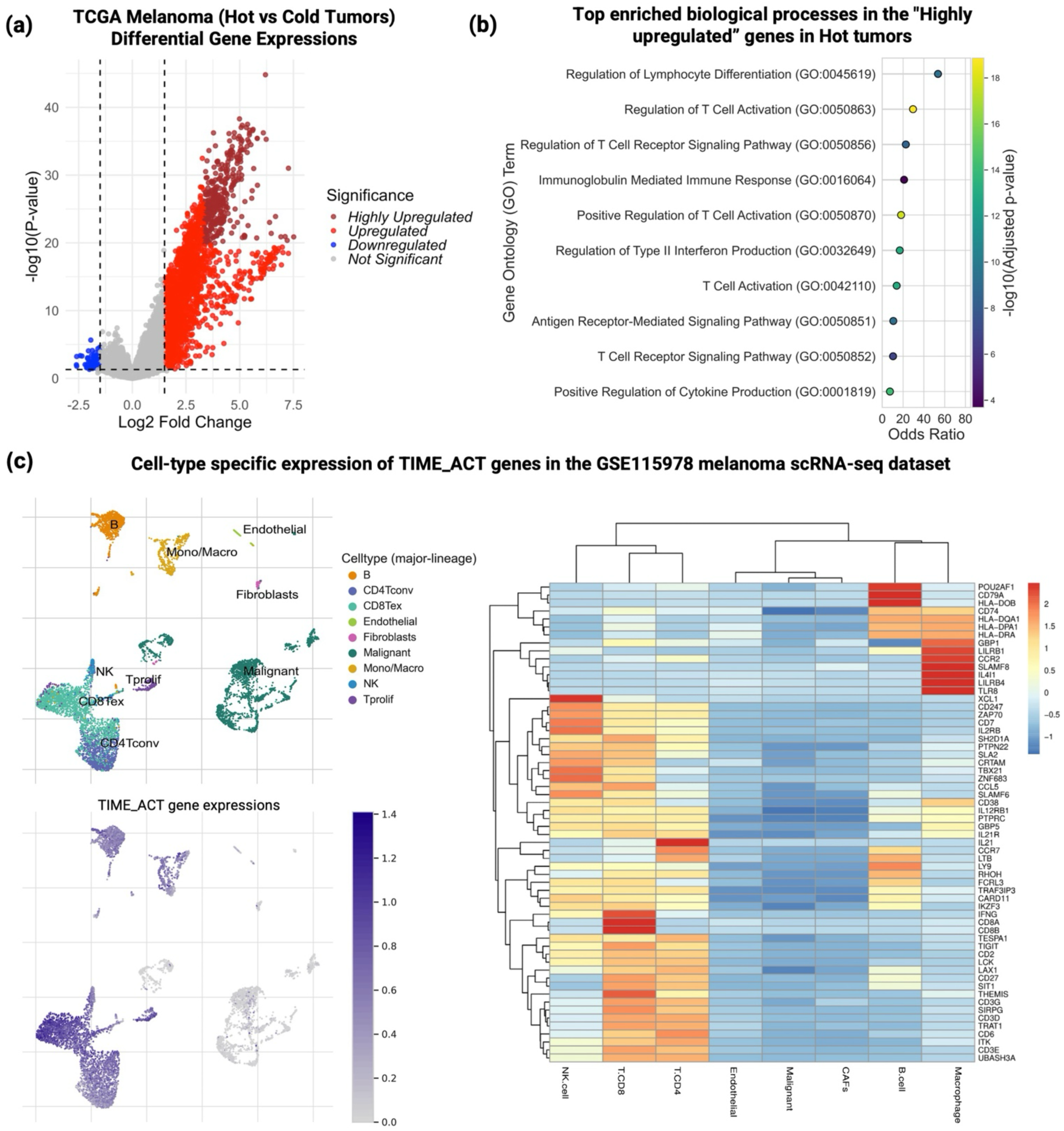
Derivation of the TIME_ACT signature by integrating immune phenotyping and transcriptomic data in melanoma. **(a)** Volcano plot showing differentially expressed genes between Hot and Cold tumors. A total of 339 genes that are upregulated in Hot vs Cold tumors were selected based on stringent thresholds for both log₂FC (>3.344) and B-statistic (>36.224). **(b)** Gene ontology enrichment analysis of the 339 upregulated genes using Enrichr. The 66 TIME_ACT genes were selected based on membership in the top ten immune-related GO Biological Process terms enriched in the 339 upregulated genes (FDR < 0.05). **(c)** Cell-type-specific expression of TIME_ACT genes in the GSE115978 single-cell RNA-seq melanoma dataset was retrieved from the TISCH2 database. The left upper panel shows UMAP clustering of transcriptionally defined cell populations. The left bottom panel overlays the same UMAP with TIME_ACT gene expression, where the intensity of purple indicates higher aggregate expression of TIME_ACT genes. The right panel represents the expression heatmap of the TIME_ACT genes (y-axis) across this dataset, with the x-axis displaying different cell types. As evident, the latter are predominantly expressed in immune cell populations, including T cells and B cells, and are minimally expressed in malignant cells and fibroblasts, supporting their immune-specific role within the TME.

To determine whether TIME_ACT reflects histopathologic immune architecture, we first evaluated its concordance with the five-tier TIL Map structural patterns (Brisk Diffuse, Brisk Band-like, Non-Brisk Multifocal, Non-Brisk Focal, and None) defined by Saltz et al ^39^. We implemented a single-sample gene set enrichment analysis (ssGSEA) using the 66-gene TIME_ACT signature to derive a TIME_ACT score for each sample (**Methods**). Across all TCGA-SKCM samples, TIME_ACT scores exhibited a strong monotonic trend, with the highest values observed in tumors displaying brisk lymphocytic infiltration and progressively lower scores in non-brisk and TIL-absent patterns (**Supplementary Fig. 1d**), indicating that transcriptomic immune activation closely mirrors the degree and structural organization of lymphocyte infiltration observed histopathologically.

To investigate whether these 66 genes represent a coherent biological program rather than a random collection of immune-associated genes, we performed Weighted Gene Co-expression Network Analysis (WGCNA) on the top 10,000 most variable genes across TCGA melanoma samples. This analysis identified multiple gene modules **(Supplementary Fig. 1e)**, among which the turquoise module emerged as the most strongly associated with the Hot phenotype (module eigengene *p = 3.69e-35*) **(Supplementary Fig. 1f)**. Importantly, all 66 TIME_ACT genes clustered within this turquoise module, underscoring their functional relatedness. To further quantify their centrality, we computed module membership scores (kME), which measure the correlation of each gene with the module eigengene. TIME_ACT genes exhibited significantly higher kME scores than other genes in the same module (*p = 1.66e-82*) **(Supplementary Fig. 1g)**, indicating that they are not only co-expressed but also represent core “hub” genes within this immune-activated transcriptional network. These results confirm that TIME_ACT genes are functionally connected and topologically central within a network specifically enriched in Hot tumors, providing strong evidence that they define a biologically meaningful immune activation signature.

### TIME_ACT scores accurately identify tumor immune activation in other cancer types

To investigate the potential applicability of the 66-gene TIME_ACT signature in other cancer types, we studied its ability to accurately classify Hot versus Cold tumors in each cancer type. We focused on eight TCGA cancer types, where corresponding immune, inflammation, and TIL patch scores were available (**Supplementary Table 4**). First, using the same top and bottom 30% strategy applied to these three scores in melanoma, tumors in each cancer type were annotated as Hot or Cold based on composite immune profiling, to generate a ‘ground truth’ dataset of annotated Hot and Cold tumors. Given that, we then tested the ability of the melanoma-derived TIME_ACT signature to distinguish these annotated groups.

Higher TIME_ACT scores indicate stronger immune activation and are characteristic of immune-hot tumors **(Fig. 3a)**. Remarkably, the TIME_ACT scores demonstrated very high classification accuracy in all cancer types, with AUC values ranging from 0.96 (for STAD) to 1.00 (BRCA and CESC) **(Fig. 3b, left)**, showing that the melanoma-derived signature accurately captures immune activation in diverse TMEs. To further test the robustness of the signature, we applied it to two independent datasets ^42^ (GSE161537-NSCLC and GSE159067-HNSCC), where gene expression data were available, and each tumor sample had been pre-annotated as Hot or Cold. Reassuringly, also in both these external cohorts, the TIME_ACT score showed excellent prediction accuracy, achieving AUC values of 0.98 (for GSE161537-NSCLC) and 0.96 (for GSE159067-HNSCC) (**Fig. 3b, right)**. These results suggest that the melanoma-derived TIME_ACT signature is not only robust across cancer types but also transferable across cohorts, platforms, and study designs.

**Fig. 3.**
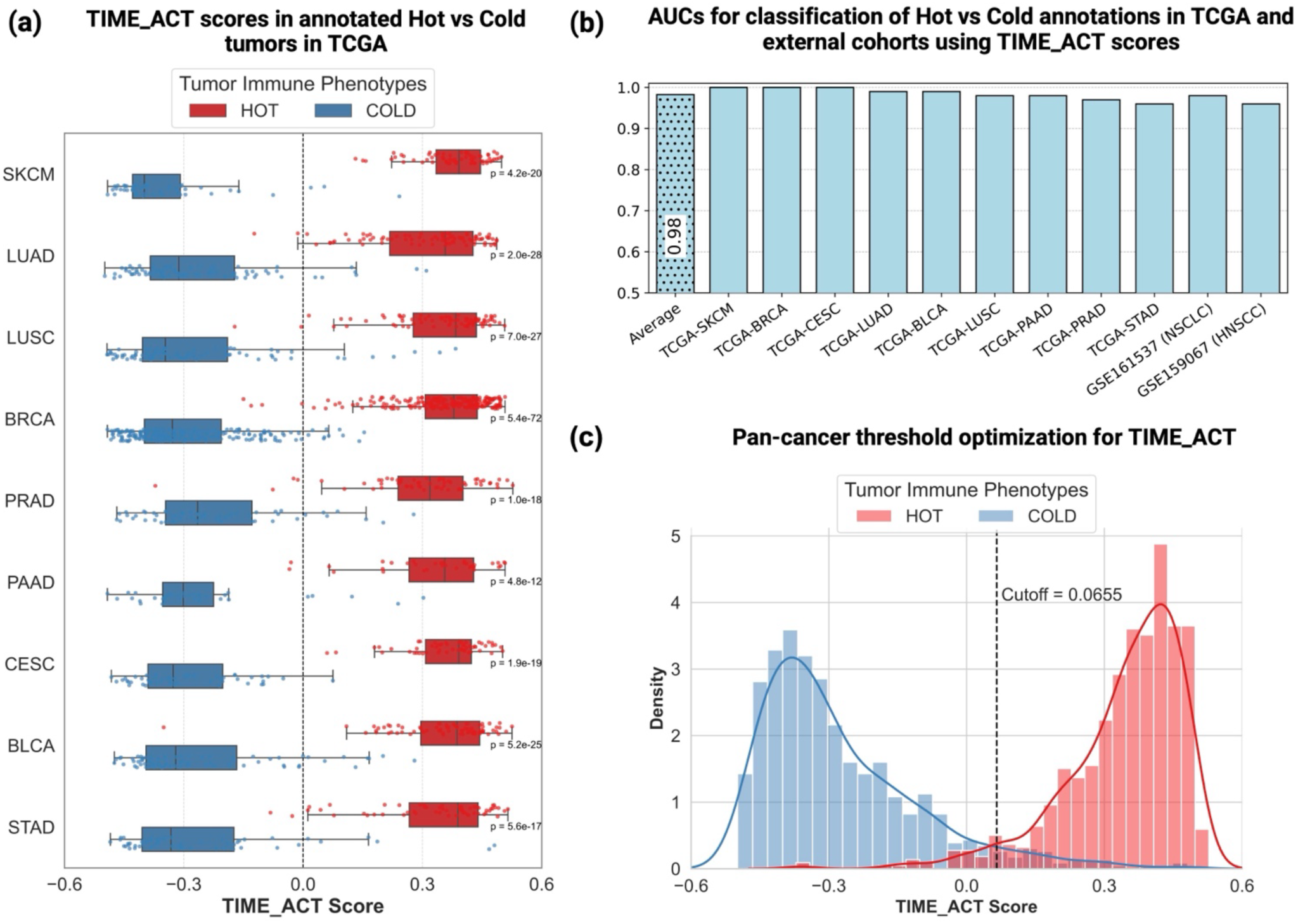
TIME_ACT defines a robust transcriptomic biomarker of tumor immune activation across cancer types. **(a)** Distribution of TIME_ACT scores in Hot and Cold annotated tumors across nine TCGA cancer types with annotated ground truth, showing consistently markedly higher scores in the originally annotated hot tumors. **(b)** Bar plot summarizing AUC of classifying Hot vs. Cold annotated tumors across the nine annotated TCGA cancer types using the TIME_ACT signature and in two independent datasets with ground truth annotated Hot/Cold labels, GSE161537 (NSCLC) and GSE159067 (HNSCC). All cohorts demonstrated strong discriminatory accuracy (AUCs ≥ 0.96). **(c)** Pan-cancer optimization of the TIME_ACT classification threshold using Youden’s index across pooled Hot and Cold samples from nine TCGA cancer types.

Next, to evaluate the fraction of Hot tumors, we computed the TIME_ACT scores for all TCGA samples **(Supplementary Table 5).** To enable consistent cross-cancer comparisons, we established a pan-cancer cut-off by pooling data from nine tumor types, including melanoma and eight additional cancers, where we annotated the Hot and Cold labels. Using the Youden Index, we identified an optimal threshold of 0.0655 that maximally separated Hot and Cold tumors based on TIME_ACT scores **(Fig. 3c)**. Notably, in all analyses shown in the following, this threshold was kept fixed and applied uniformly across the TCGA and any other patients’ datasets analyzed herewith.

### Studying the biological bases of the TIME_ACT signature at bulk, single-cell, and spatial resolutions

To further contextualize the biological basis of TIME_ACT and assess its cross-tumor applicability, we performed an integrative, pan-cancer analysis leveraging bulk transcriptomics, immune subtyping, single-cell datasets, and spatial data across diverse cancer types. First, we obtained the immune subtype annotations and cell-type abundance estimates from the TCGA immune landscape resource described by Thorsson et al. ^43^, which used pan-immune gene expression features and deconvolution-based immune cell estimates and classified tumors into 6 different immune subtypes (C1–C6). To test whether TIME_ACT scores tracked with these established immune-active phenotypes, we examined the distribution of immune subtypes in TIME_ACT-defined Hot tumors. The majority of Hot tumors fell into immune-active subtypes, either the C2 (IFN-γ dominant; 40.1%) or C3 (inflammatory; 30.2%) **(Fig. 4a)**, both of which are associated with elevated cytotoxicity, antigen presentation, and lymphocyte infiltration. Consistent with this, TIME_ACT scores showed strong correlations with key pan-cancer immune signatures, including IFN-γ response, lymphocyte infiltration, TCR and BCR richness, and leukocyte fraction **(Fig. 4b)**. Together, these findings further signify that the TIME_ACT signature reliably captures tumors with immune-inflamed phenotypes characterized by active antitumor immune responses across cancer types.

**Fig. 4.**
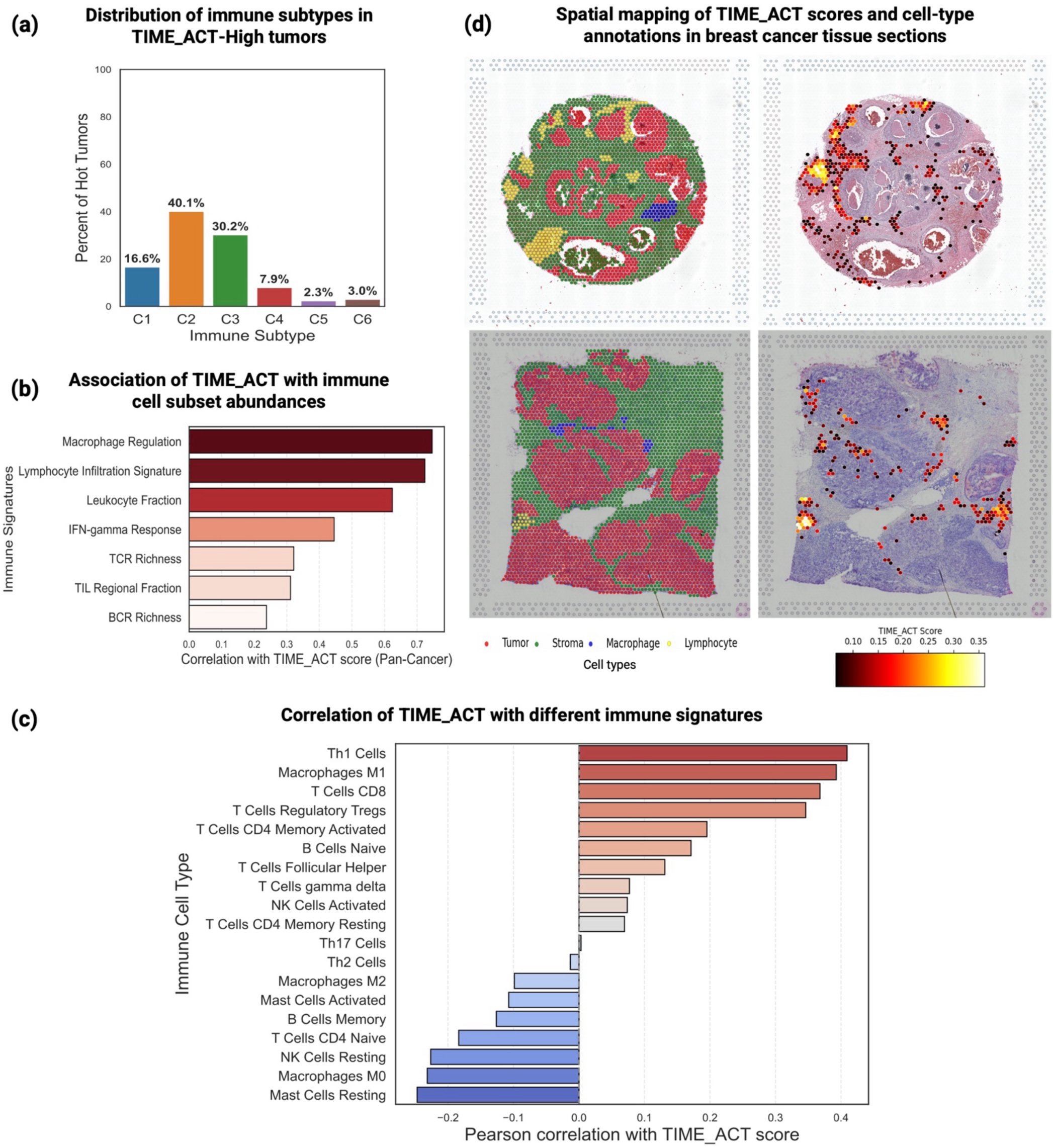
Multiscale characterization of TIME_ACT across immune subtypes, cellular composition, and spatial localization in diverse cancer types. **(a)** Distribution of immune subtypes among TIME_ACT-defined Hot tumors across the TCGA pan-cancer cohort. Hot tumors are predominantly classified as C2 (IFN-γ dominant) and C3 (inflammatory) subtypes, known for immune-inflamed phenotypes. **(b)** Correlation of TIME_ACT scores with established immune-related signatures across cancers, including IFN-γ response, lymphocyte infiltration, TCR/BCR richness, and leukocyte fraction, demonstrating alignment with active immune microenvironments. **(c)** Association of TIME_ACT scores with immune cell subset abundance from deconvolved TCGA data. Positive correlations were observed with CD8⁺ T cells, M1 macrophages, and Th1 cells; negative correlations with resting immune populations such as M0 macrophages and resting NK cells. **(d)** Spatial transcriptomics analysis of breast cancer tissue sections from the 10x Genomics repository shows that TIME_ACT-high (hot) regions are enriched in lymphocyte-dense areas adjacent to tumor epithelium.

We then tested whether TIME_ACT genes consistently localize to immune compartments at the single-cell level across tumor types, extending our previous analysis in melanoma. Consistently, we find that TIME_ACT genes remained preferentially expressed in immune cells, particularly T cells and B cells, and were rarely expressed in malignant or stromal compartments **(Supplementary Fig. 2a)**. Given this strong immune-cell specificity, we next asked whether TIME_ACT scores correlate with the abundance of immune cell types known to mediate antitumor immunity, using deconvolved bulk transcriptomic data from TCGA. Across cancer types, TIME_ACT was positively correlated with CD8⁺ T cells, M1 macrophages, and Th1 cells, immune subsets canonically associated with pro-inflammatory and cytotoxic activity **(Fig. 4c)**. In contrast, TIME_ACT scores were negatively correlated with resting or undifferentiated populations, including M0 macrophages, resting NK cells, and resting mast cells. These associations suggest that higher TIME_ACT scores are linked to transcriptional programs characteristic of pro-inflammatory and cytotoxic immune cell states.

Finally, we studied what key spatial relations between different cell-types in the TME may be associated with immune activation as quantified by TIME_ACT scores. To this end, we analyzed breast cancer tissue sections, both formalin-fixed paraffin-embedded and fresh-frozen, from the 10x Genomics public spatial transcriptomics repository. Each spatial spot in this cohort was annotated by expert pathologists to classify tumor cells, stromal cells, lymphocytes, and macrophages. TIME_ACT scores were computed per spot, and those exceeding the pan-cancer threshold (≥ 0.0655), were designated as Hot. In both samples, we observed that high TIME_ACT-scoring regions tended to co-localize with spots enriched in dense lymphocyte aggregations that are adjacent to malignant areas **(Fig. 4d)**. This analysis testifies to a spatial coherence between transcriptomically inferred immune activation and histologically defined lymphocyte localization in the TME. Together, these findings demonstrate that TIME_ACT reflects not only molecular programs of immune activation but also can be applied to uncover meaningful spatial relationships between immune and malignant cells, reinforcing its role as a multiscale biomarker of inflamed TMEs.

### TIME_ACT outperforms numerous published transcriptomic-based signatures and methods of ICB response prediction

We have seen that TIME_ACT scores can distinguish between hot and cold tumors within the same cancer type. To assess whether TIME_ACT scores in tumors with confirmed immune activation associate with clinical benefit from ICB, we analyzed objective response rates (ORRs) to ICB therapy across nine cancer types. Specifically, we studied whether higher mean TIME_ACT scores among these Hot tumors correlate with higher ORRs across cancer types. ORR data were retrieved from our previous studies ^44,45^. Among the nine cancer types evaluated, eight had both ORR and TIME_ACT data available. Remarkably, we observed a strong positive correlation between the mean TIME_ACT score of Hot tumors and the corresponding ORR (Pearson r = 0.72, *p = 4.46e-02*; **Supplementary Fig. 3a**), suggesting that tumors with more pronounced immune activation, as captured by TIME_ACT, are more likely to respond to ICB therapy. These findings support the translational relevance of the TIME_ACT signature as a pan-cancer biomarker of immune activation and ICB response, setting the stage for the next steps, where we explore the predictive power of TIME_ACT scores of individual patient’s tumors to predict their response to ICB therapy.

To this end, we evaluated TIME_ACT in pretreatment tumor transcriptomes from 25 independent ICB-treated cohorts spanning nine cancer types and several treatment settings, including anti-PD-1, anti-CTLA-4 and combination therapy **(Supplementary Table 6)**. None of these cohorts was used to derive the TIME_ACT signature, thereby providing an independent assessment of its generalizability. TIME_ACT predicted response across diverse cancer types and treatment contexts **(Supplementary Fig. 3b)**, with a mean area under the receiver operating characteristic curve (AUC) of 0.76 across the 25 cohorts. Cancer type-specific mean AUCs were 0.76 across six melanoma cohorts, 0.83 across three non-small-cell lung cancer (NSCLC) cohorts, 0.74 across four breast cancer cohorts, 0.70 across five clear-cell renal cell carcinoma (ccRCC) cohorts and 0.78 across two head and neck squamous cell carcinoma (HNSCC) cohorts. Predictive performance was also observed in individual cohorts of mesothelioma, pancreatic cancer, prostate cancer and sarcoma, supporting the applicability of TIME_ACT beyond the more extensively represented indications **(Supplementary Fig. 3c, upper panel)**.

We next evaluated whether a single, prespecified TIME_ACT threshold could possibly provide a translationally useful response prediction for patients, by maximizing the Odds Ratios (OR) of true responders among those predicted to response (with scores greater than that threshold) vs those predicted to be non-responders. The previously established pan-cancer threshold of 0.0655 was applied as is to stratify the patients in each cohort to responder’s vs non-responders, accordingly. Across the 25 cohorts, TIME_ACT-high tumors were enriched for responders, with a mean odds ratio (OR) of 5.77. Cancer type-specific mean ORs were 6.89 for melanoma, 4.44 for NSCLC, 6.43 for breast cancer, 3.86 for ccRCC and 8.77 for HNSCC (**Supplementary Fig. 3d, lower panel**). Consistent with the threshold-based analysis, responders generally had higher continuous TIME_ACT scores than non-responders across the 25 cohorts (**Supplementary Fig. 4**), further reinforcing the signature’s strong association with clinical benefit. These findings emphasize the translational potential of TIME_ACT not only as a continuous signature but also as a threshold-based, clinically actionable predictor of response to ICB therapy across diverse tumor types.

We then benchmarked TIME_ACT against 21 published immune-related gene-expression signatures across all 25 pretreatment cohorts **(Supplementary Table 7)**. The comparator panel represented major biological components previously associated with ICB response, including IFN-γ signaling, cytolytic activity, T- and B-cell programs, macrophage and myeloid-cell states, antigen-presentation machinery, immune-checkpoint expression and TGF-β signaling. The analysis also included the clinically relevant single-gene markers PDCD1 (PD-1), CD274 (PD-L1) and CTLA4. TIME_ACT achieved a significantly higher mean AUC than every comparator signature and had the lowest coefficient of variation (CV) across cohorts, indicating that its performance was not driven by a small number of favorable datasets but was comparatively stable across cancer types, treatment regimens and study populations **(Fig. 5 a, b)**.

**Fig. 5.**
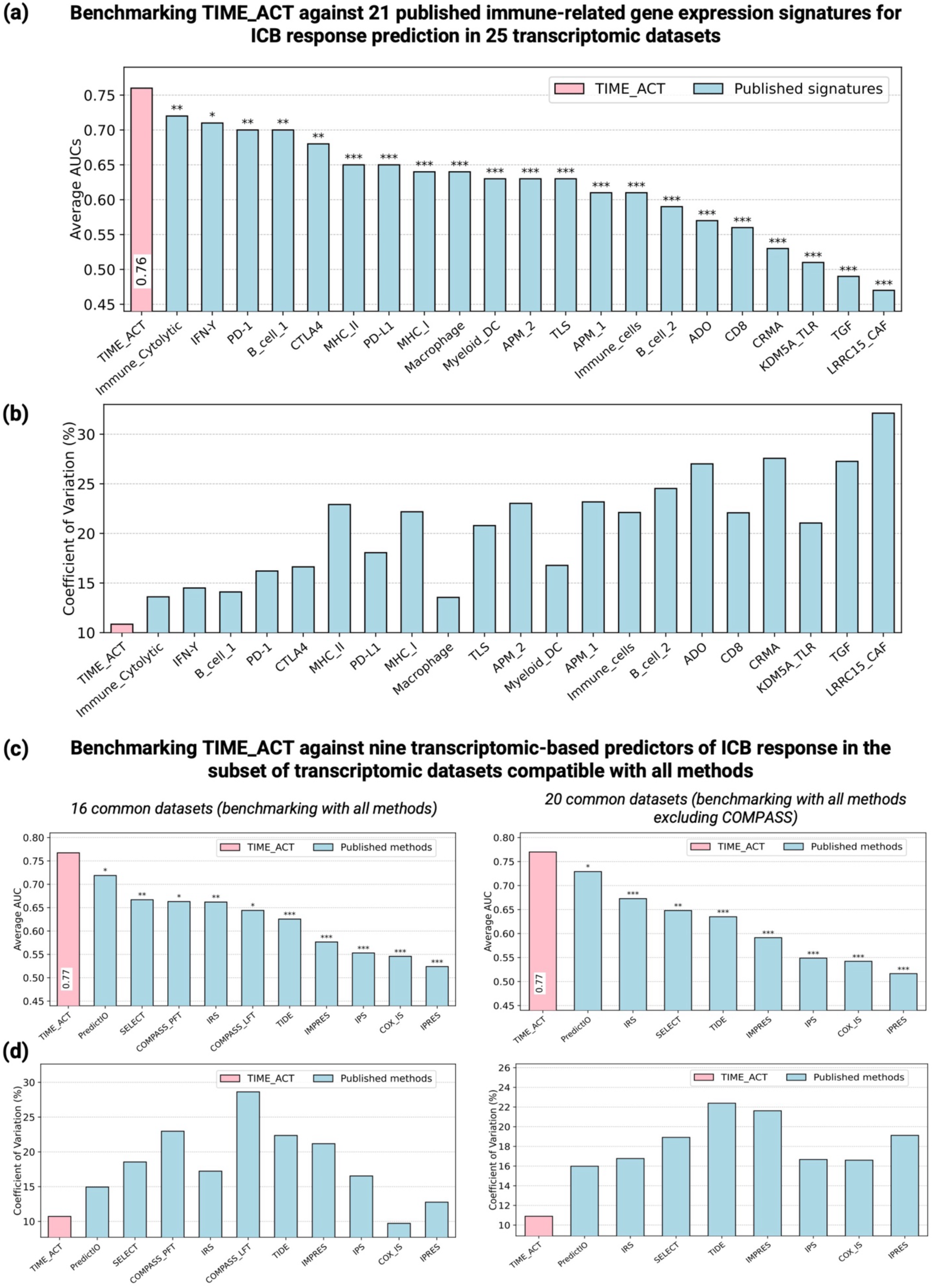
Benchmarking TIME_ACT against published immune-related ICB response gene signatures and transcriptomic-based prediction methods. **(a-b)** Comparative performance of TIME_ACT versus 21 published immune-related gene expression signatures for ICB response prediction across 25 independent pre-treatment cohorts. Bars indicate mean AUC (top) and corresponding coefficient of variation (CV, bottom) across all cohorts. TIME_ACT achieved the highest mean AUC and the lowest inter-cohort variability. **(c-d)** Head-to-head benchmarking of TIME_ACT against nine transcriptomic-based prediction methods using matched cohorts. The left panel shows the strictly matched comparison across 16 cohorts in which all methods, including COMPASS-PFT and COMPASS-LFT, could be evaluated. The right panel shows the comparison across 20 common cohorts shared by TIME_ACT and all remaining methods after excluding COMPASS, which could not be reliably applied to additional cohorts because of insufficient coverage of its required input genes. Bars indicate mean AUC and **(d)** Corresponding CVs for the 16-cohort and 20-cohort matched comparisons. TIME_ACT achieved the highest mean AUC and the lowest cross-cohort variability in both analyses. Statistical significance was assessed using one-sided paired Wilcoxon signed-rank tests comparing cohort-level AUCs for TIME_ACT with each comparator. *P < 0.05 (*)*, *P < 0.01 (**)* and *P < 0.001 (***)*.

Moving from signatures to compare TIME_ACT with established transcriptome-based ICB response predictor models, we systematically compared its performance with PredictIO, TIDE, SELECT, IRS, IMPRES, IPS, Cox-IS and IPRES, together with the recently developed COMPASS foundation model ^34^. We evaluated both COMPASS partial fine-tuning (PFT) and linear-probing (LFT) configurations, which were reported as its strongest and most stable implementations across clinical cohorts. Several cohorts had insufficient coverage of the genes required by COMPASS and could not be evaluated using these models. Consequently, the fully matched head-to-head comparison of TIME_ACT, COMPASS and all other methods was restricted to 16 common cohorts where it was fully feasible. TIME_ACT achieved a mean AUC of 0.77, markedly and significantly exceeding both COMPASS-PFT (AUC of 0.66) and COMPASS-LFT (AUC of 0.64), as well as the other transcriptomic predictors **(Fig. 5c, left)**. TIME_ACT also showed the lowest CV, supporting greater consistency across the matched cohorts **(Fig. 5d, left)**. As most of the remaining methods could be applied to a broader set of datasets, we performed a second matched comparison after excluding COMPASS. This analysis included 20 cohorts shared by TIME_ACT and all remaining methods. TIME_ACT again achieved the highest mean AUC, at 0.77, and the lowest CV **(Fig. 5 c,d, right)**. Together, the 16-cohort analysis provides a direct comparison with the recent COMPASS foundation model, whereas the 20-cohort analysis extends the benchmarking to the largest common set available for the remaining transcriptomic predictors.

We next assessed performance using the area under the precision-recall curve (AUPRC), a complementary metric that is particularly informative when the numbers of responders and non-responders are imbalanced. Across the 25 cohorts, TIME_ACT achieved a mean AUPRC of 0.61, exceeding those of the evaluated gene-expression signatures, and outperformed all nine previously published prediction methods across the 16 common datasets, achieving a mean AUPRC of 0.66. **(Supplementary Fig. 5 a-c)**. Thus, TIME_ACT more effectively prioritized true responders while maintaining robust performance across cohorts with different response prevalences.

We further studied whether TIME_ACT could capture not only baseline immune states but also the evolving antitumor immune response after treatment initiation, we next examined cohorts with paired pretreatment and on-treatment transcriptomic data. In the two melanoma anti-PD1 cohorts with both pre-treatment and on-treatment samples (GSE91061 and PRJEB23709) ^46,47^, TIME_ACT scores achieved AUCs of 0.66 (for GSE91061) and 0.85 (for PRJEB2370_PD1) based on the pre-treatment transcriptomics data. However, interestingly, its performance improved markedly in the corresponding on-treatment samples, reaching 0.75 (for GSE91061) and 0.93 (for PRJEB23709_PD1). These results demonstrate that TIME_ACT not only reflects pre-existing immune activity but also sensitively captures early immunologic changes induced by anti-PD1 therapy.

Finally, given the widespread clinical use of tumor mutational burden (TMB) as a biomarker for immunotherapy response, we sought to additionally benchmark TIME_ACT against TMB in a subset of patients. However, among all 25 ICB cohorts, only the Riaz et al. study ^47^ included both transcriptomic data and the necessary information for TMB calculation. Using this data, we found that TIME_ACT outperformed TMB in predicting ICB response, achieving an AUC of 0.66 versus 0.57 **(Supplementary Fig. 5d)**. Notably, TIME_ACT maintained predictive accuracy across both TMB-High (AUC = 0.64) and TMB-Low (AUC = 0.68) subgroups, underscoring its potential utility in TMB-low tumors where genomic biomarkers often underperform **(Supplementary Fig. 5e)**.

### Predicting TIME_ACT scores and treatment response directly from histopathology

Tumor histopathology remains the most accessible and widely used diagnostic modality in oncology. Building on the superior performance of TIME_ACT in predicting treatment response from transcriptomic data, we explored the feasibility of inferring the expression of the 66 genes composing TIME_ACT and based on that, computing TIME_ACT scores directly from pathology slides. Having these inferred scores, we then aimed to predict treatment response (**Fig. 1c**).

To this end, we first employed our recently published Path2Omics framework ^36^, a deep learning model that extends the capabilities of the previously established DeepPT approach ^35^, to infer gene expression from pathology slides. Notably, for predicting ICB response from slides, traditional “direct” supervised approaches predict response directly from slides without an intermediate gene expression inference step. However, these approaches typically require large numbers of samples with matched slides and response labels for each indication and treatment regime. Our “indirect” approach addresses this data limitation by decomposing prediction into two interpretable steps. First, we leverage large, publicly available TCGA resources (with over 20,000 matched slides and gene expression profiles) to train models that infer gene expression from slides. After training, the pre-trained model is used to predict gene expression from slides in the clinical treatment cohort, without any further re-training. Second, the indirect approach enables one to readily apply other validated molecular signatures to predict treatment response using the inferred gene expression, in comparison. The TIME_ACT score is computed directly from inferred gene expression, in an unsupervised learning manner, without training on the treatment cohort. Beyond the improved performance, the intermediate gene expression step makes our “indirect” approach more interpretable and has value on its own, a desired property for enhancing clinical deployment. This design improves robustness, interpretability, and cross-cohort generalizability compared to direct image-only models. Path2Omics was trained on all available TCGA slides for each cancer type, excluding those of low quality (damaged, blurry, out of focus). To avoid information leakage, the 5-fold cross-validation split was performed at the patient level, ensuring that slides from the same patient were assigned to the same fold. Predictions at the patient level were then obtained by averaging the predictions made at the slide level. Path2Omics was first trained and cross-validated within the TCGA cohort and then externally validated on seven independent cohorts. To assess the inference accuracy, we calculated the Pearson correlation between inferred and actual expression levels for each of the TIME_ACT genes across all test samples within each cancer type. Notably, the inferred gene expression values were obtained exclusively from test sets that remained unseen during Path2Omics training, mitigating concerns about overfitting. Through fivefold nested cross-validation within each cancer type in the TCGA cohort, Path2Omics demonstrated strong performance for the 66 TIME_ACT genes, with a median correlation of 0.60 on average across 22 major cancer types, ranging from 0.42 (for PCPG) to 0.71 (for TGCT) (**Fig. 6a**, **pink**). Notably, this performance was remarkably higher than that observed for all genes (**Fig. 6a, light blue**), indicating that the TIME_ACT genes are among the most accurately inferred by Path2Omics. We further evaluated Path2Omics in seven external validation cohorts comprising 1,229 samples across six cancer types and observed a consistent performance trend. Path2Omics achieved a median correlation of 0.56 on average across these external cohorts, again outperforming its performance on all genes (**Fig. 6b**). These findings underscore the robustness and generalizability of Path2Omics.

**Fig. 6.**
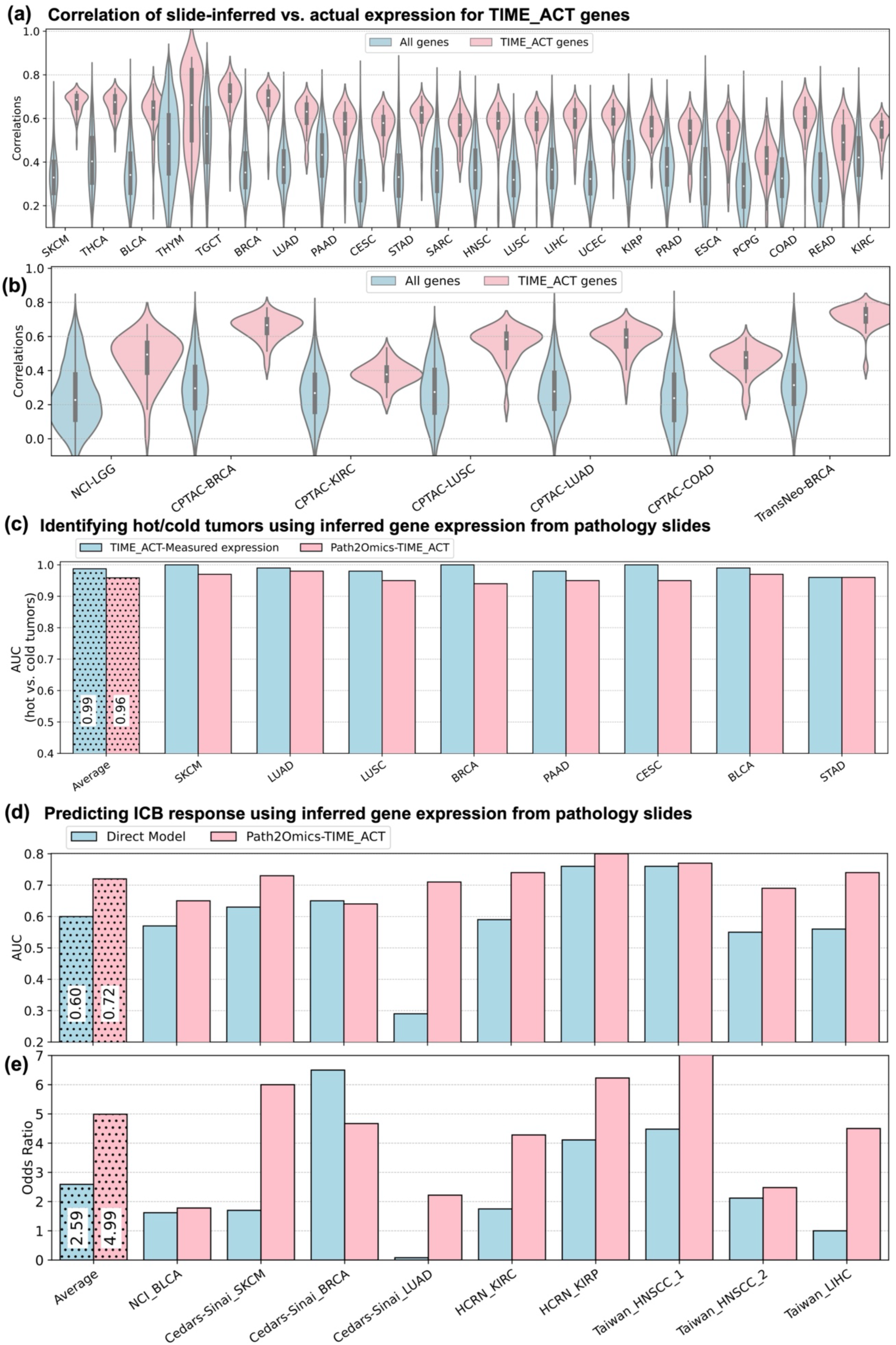
Predicting expression of TIME_ACT genes and treatment response from pathology slides. **(a)** Violin plots demonstrate the distribution of Pearson correlation coefficients between slide-inferred and measured RNA-seq expression for TIME_ACT genes (orange) versus all genes (gray) in different TCGA cancer types. **(b)** Boxplots summarize the performance of Path2Omics across seven independent external validation cohorts spanning six cancer types (n = 1,229 samples). Performance is measured as the Pearson correlation between slide-inferred and matched RNA-seq expression. **(c)** Bar plots show performance in predicting Hot vs. Cold tumors in different cancer types, based on slide-inferred TIME_ACT scores (orange) and TIME_ACT measured expression (blue). AUC values for slide-inferred scores closely match those from RNA-seq-based scores. **(d)** Comparison of AUC values between slide-inferred TIME_ACT scores and a direct histopathology-based model for predicting ICB response across nine independent ICB-treated clinical cohorts. **(e)** Comparison of odds ratio (OR) values between slide-inferred TIME_ACT scores and a direct histopathology-based model trained to predict ICB response. Across all cohorts, TIME_ACT outperforms the direct model (mean AUC 0.72 vs. 0.60; mean OR 4.99 vs. 2.59), underscoring the advantage of transcriptional imputation prior to response prediction.

We then computed TIME_ACT scores based on the slide-inferred gene expression and evaluated their ability to classify tumors into “Hot” and “Cold” categories, analyzing the TCGA cancer types with the ground truth Hot/Cold annotations. Remarkably, the slide-inferred TIME_ACT scores achieved AUC values exceeding 0.94 in all tested cancer types, closely matching the performance of TIME_ACT scores computed from actual measured expression data **(Fig. 6c)**. Notably, the results were obtained from test sets that remained unseen during the training of Path2Omics, alleviating potential concerns about overfitting. Taken together, these findings demonstrate that TIME_ACT gene expression and composite scores can be reliably inferred from pathology slides to a level that is sufficient for detecting tumor hotness directly from the tumor pathology slides.

We next applied the TCGA cancer type-specific pre-trained Path2Omics model to predict gene expression and subsequently computed the TIME_ACT scores of each patient directly from their tumor histopathology slides in a collection of 9 multi-centric ICB clinical cohorts for which we had the slides and response data. Those include nine unpublished cohorts of patients treated with ICB, from multiple different medical centers, including datasets from our in-house NCI, Cedars-Sinai, HCRN (Hoosier Cancer Research Network) consortium, and NYCU-Taiwan. Notably, ICB response prediction based on the slide-inferred TIME_ACT scores achieved a mean AUC of 0.72, ranging from 0.64 (Cedars-Sinai BRCA cohort) to 0.8 (HCRN KIRP cohort) (**Fig. 6d**). The mean OR across these ICB datasets was 4.99, ranging from 1.78 (NCI BLCA cohort) to 12.75 (NYCU-Taiwan HNSCC cohort) (**Fig. 6e**). Notably, this performance was markedly higher than that achieved by a traditional “direct” modeling approach, which predicts treatment response directly from slides, without an intermediate gene expression inference step, achieving a mean AUC of 0.60 (**Fig. 6d**, left, in blue) and mean OR of 2.59 (**Fig. 6e**, left, in blue). These findings indicate that incorporating biologically interpretable transcriptional information enhances both the accuracy and robustness of slide-based predictors. The superiority of the “indirect” approach over the “direct” approach has also been consistently observed in our previous studies ^35,36^.

The ROC curves for each individual cohort and the distribution of slide-inferred TIME_ACT scores between responders and non-responders across cohorts are shown in **Supplementary Fig. 6a** and **Supplementary Fig. 6b**, respectively. Among all the slide-based cohorts we analyzed, matched RNA-seq data were available for the HCRN-KIRC cohort (dbGaP accession: phs003618.v1.p1). This enabled us to evaluate Path2Omics performance on this cohort. Once again, we found that TIME_ACT genes are among the well-predicted genes via Path2Omics, with a median correlation of 0.49, remarkably outperformed the median correlation of 0.18 for all genes (**Supplementary Fig. 6c**). This dataset also enabled us to perform a direct comparison of the predictive performance of TIME_ACT scores derived from slide-inferred expression versus those computed from measured RNA-seq data. Remarkably, the slide-inferred TIME_ACT scores, based on histology-derived expression, matched the performance of RNA-seq-based scores. In the subset of matched HCRN-KIRC cohort (n = 82), TIME_ACT scores based on slide-inferred expression achieved an AUC of 0.76 and OR of 5.96, as good as the scores derived based on RNA-seq-based expression data (AUC of 0.72 and OR of 4.52). Taken together, these results underscore the predictive capability of the combined TIME_ACT and Path2Omics in predicting patient response directly from tumor pathology slides, offering a rapid and cost-effective alternative to traditional precision oncology omics-based tools.

## Discussion

Accurately predicting a patient’s response to ICB remains an urgent unmet clinical need, as it is crucial for guiding optimal treatment decisions ^1^. However, the existing transcriptomic biomarkers for ICB response often suffer from unstable performance on unseen datasets, limiting their clinical applicability ^32,33,48^. This study presents TIME_ACT, a transcriptomic signature of tumor immune activation, which offers a simple, unsupervised approach for identifying immune-hot tumors and predicting therapeutic response to ICB treatments in many different cancer indications. Notably, the robust predictive power of TIME_ACT extends to digital pathology, maintaining a strong predictive power across diverse clinical settings. Using spatial transcriptomics and digital histopathology, we show that TIME_ACT-high regions co-localize with immune effector cells in direct proximity to tumor cells, reinforcing that the signature captures key spatial features of an inflamed TME.

The ability of TIME_ACT to generalize across cohorts and data modalities is likely rooted in its representation of tumor hotness as a multidimensional biological state rather than as a single immune feature. TIME_ACT is strongly associated with several established features of an inflamed tumor microenvironment, including IFN-γ response, lymphocyte infiltration, leukocyte fraction, T-cell receptor richness and B-cell receptor richness. It also correlated with immune-cell states linked to effective antitumor immunity, including CD8⁺ T cells, Th1 cells and M1 macrophages, while showing inverse associations with resting or less differentiated immune populations. This broader integrative biological coverage may explain why TIME_ACT significantly outperformed all 21 evaluated immune-related gene-expression signatures for ICB response prediction. Each of the latter covers important but specific and narrow features determining the effectiveness of the immune response, whose predictive value can vary across cancer types. By integrating multiple dimensions of tumor hotness, TIME_ACT provides a broader and more stable representation of the immune microenvironment across cohorts. Indeed, TIME_ACT predictive power has the lowest coefficient of variation across cohorts, testifying to its robustness.

The benchmarking against established transcriptomic prediction methods further supports the value of this biologically constrained representation. Particularly, the comparison with COMPASS provides an important contemporary benchmark. COMPASS is a recently developed pan-cancer foundation model that combines a transformer-based gene encoder, biologically defined immune concepts and pretraining on more than 10,000 tumors, followed by cohort-specific adaptation. In the 16 transcriptomic cohorts that could be evaluated by all approaches, TIME_ACT significantly outperformed both the PFT and LFT COMPASS configurations. This result is particularly notable because TIME_ACT is a fixed, unsupervised gene-expression signature that requires neither model training nor cohort-specific fine-tuning, yet it outperformed a substantially more complex foundation model pretrained on more than 10,000 tumors. This indicates that, for quantifying a dominant and biologically coherent axis such as tumor immune activation, a carefully derived signature can retain substantial predictive information without the additional complexity of large-scale model training.

Importantly, the performance of TIME_ACT extended to new clinical cohorts where TIME_ACT scores were inferred directly from histopathology slides, achieving predictive accuracy comparable to RNA-seq-derived scores. This modality-bridging capability distinguishes TIME_ACT from existing transcriptomic signatures and direct image-only predictors and establishes a practical foundation for a future clinically deployable immune activation scoring across cancer types, upon further careful prospective testing. If successful, this may hold promise for advancing precision ICB treatment in developing and resource-constrained healthcare systems, with limited access to comprehensive molecular profiling.

This study has a few limitations. First, our analysis was retrospective in nature, and while the TIME_ACT score showed strong predictive performance across multiple cohorts and modalities, including both RNA-seq and slide-inferred expression data, prospective validation in clinical settings is still needed. Second, while the current framework focuses on transcriptome-derived features, it does not incorporate histological or spatial characteristics of the tumor microenvironment. With the emergence of slide-based gene expression inference, future efforts may explore integrating TIME_ACT with spatially resolved features to capture localized immune deserts, excluded regions, or inflamed niches that may be missed by conventional bulk RNA-seq ^49^. Such integration could enhance the precision and applicability of TIME_ACT for guiding immunotherapy decisions in the clinic. Third, we acknowledge that immune ‘hotness’ represents only one component of ICB response biology. Many tumors possess distinct immunological features, including myeloid-driven immunosuppression, angiogenesis-mediated immune exclusion, and metabolic reprogramming, which may modulate therapeutic response independently of lymphocyte infiltration. Accordingly, TIME_ACT is designed to quantify tumor immune activation rather than to fully capture all determinants of ICB responsiveness across cancer types. Integrating tumor immune activation with additional cancer-specific immune axes represents an important direction for future work.

In summary, TIME_ACT is a biologically coherent, unsupervised transcriptomic signature biomarker of tumor immune activation, which robustly predicts ICB response across diverse tumor types via histology, thus offering an exciting potential for broad clinical utility. It thus lays a solid basis for further prospective studies to test its future use in the clinic as a first-of-its-kind, fast, cost-effective, and reliable ICB treatment biomarker.

## Methods

### Transcriptomic data acquisition and processing of raw RNA-seq data

Gene-expression and clinical data for The Cancer Genome Atlas (TCGA) and TARGET projects were obtained from the UCSC Xena portal ^50^ (last accessed 4 January 2025). Pre-processed STAR-aligned RNA-seq data (log₂[count + 1]) for tumor samples in the TCGA Skin Cutaneous Melanoma (TCGA-SKCM) cohort were downloaded from the UCSC Xena data portal. For pan-cancer analyses, we retrieved RSEM-based, normalized RNA-seq data (log₂[norm_count + 1]) for tumor samples from the TCGA + TARGET Pan-Cancer (PANCAN) cohort.

We selected publicly available ICB-treated cohorts only if gene expression data were generated using RNA-seq experiments and clinical response annotations were available. Microarray-based datasets were excluded due to limited gene coverage and platform-specific biases that could compromise cross-cohort comparability. We have collected 25 ICB pre-treatment (total 1196 patient samples), and 2 anti-PD1 on-treatment (total 74 patient samples). Clinical response was uniformly defined using RECIST criteria ^51^, with complete response (CR) and partial response (PR) classified as responders, and stable disease (SD) and progressive disease (PD) as non-responders. To benchmark predictive performance, we compared TIME_ACT with other transcriptomic-based methods and immune-related ICB response gene signatures in 25 ICB datasets.

Gene-level TPM expression data were obtained where available directly from the GEO database or from supplementary materials provided by the respective publications. Several datasets were dbGaP-controlled, for which authorized access was obtained. Raw FASTQ files for these restricted-access cohorts were downloaded using the fasterq-dump utility (SRA Toolkit) with the provided dbGaP keys. Transcript abundance was quantified using Salmon ^52^ against the GENCODE GRCh38 reference transcriptome. Gene-level TPM values were then derived from transcript-level estimates using the tximport R package ^53^. All TPM values were transformed to log₂(TPM + 1) prior to analysis. Single-cell gene expression data of melanoma patients were collected from the GSE115978 dataset ^40^.

### Immune scoring-based classification of TCGA tumor samples

To assess the immune landscape of tumor samples, we utilized three independent immunophenotyping metrics: immune score, inflammatory response score, and number of tumor-infiltrating lymphocyte (TIL) patches. The Immune score of the TCGA samples was derived using the ESTIMATE algorithm ^37^. The inflammatory response score of the TCGA samples was obtained from the Supplementary Data of Chen et al ^38^. The Number of TIL patches per tumor sample was collected from Saltz et al. ^39^, which applied a deep learning-based image analysis pipeline to standard H&E-stained whole-slide images (WSIs) across the TCGA cohort to identify and quantify TIL-rich regions. These values were obtained directly from the publicly shared resource linked to the publication. For each metric, samples were ranked and dichotomized into immune-high (top 30%) and immune-low (bottom 30%) groups. To ensure stringent and biologically consistent classification, we defined “immune-active” (or “Hot”) tumors as those that fell within the top 30% for at least two out of the three scores and in none of the bottom 30%. Conversely, “immune-inactive” (or “Cold”) tumors were those in the bottom 30% for at least two scores and not in the top 30% for any. Samples not meeting these criteria were excluded from downstream binary comparisons to avoid intermediate or ambiguous immune phenotypes. This strict consensus-based approach enabled the selection of only the true Hot and Cold tumors, serving as a high-confidence foundation for identifying a transcriptional signature of tumor immune activation or tumor hotness.

### Differential expression analysis, pathway analysis, and derivation of TIME_ACT signature genes

To identify genes associated with tumor immune activation, we performed differential gene expression analysis using the limma package ^54^ between Hot (n = 52) and Cold (n = 45) tumors in the TCGA Skin Cutaneous Melanoma (TCGA-SKCM) cohort, as defined by the consensus-based immunophenotyping criteria described in the previous section. Genes were ranked by both moderated log₂ fold change and B-statistic, the latter representing the log-odds of true differential expression under empirical Bayes moderation. Genes falling within the top quartile for both metrics (log₂FC > 3.344 and B > 36.224) were considered highly upregulated, resulting in a candidate set of 339 genes. Functional enrichment of these genes was conducted using the Enrichr platform ^55^, focusing on Gene Ontology (GO) Biological Process annotations. Significantly enriched pathways were identified based on adjusted p-values (Benjamini-Hochberg FDR < 0.05). Among the top 10 ranked GO terms, all were related to immune processes, including lymphocyte activation, T cell differentiation, and antigen presentation. Genes associated with these top ten immune-related categories were aggregated, yielding a final set of 66 genes, herein referred to as the TIME_ACT signature.

### Weighted gene-co-expression network analysis

We applied WGCNA v1.73 ^56^ to the top 10,000 most variable genes in the TCGA-SKCM cohort to identify co-expression modules associated with the immune-hot phenotype. Samples were hierarchically clustered to detect outliers (cut height = 100), with none removed. A soft-thresholding power of 6 was selected based on the scale-free topology criterion (R² > 0.8). Using an unsigned topological overlap matrix (TOM), modules were detected via dynamic tree cutting. Module eigengenes were correlated with the Hot vs. Cold phenotype using Pearson correlation; the turquoise module showed the strongest positive association with immune-hot tumors. Gene centrality was quantified using module membership (kME), defined as the correlation between gene expression and the module eigengene. This allowed us to assess the topological importance of genes within each module and compare the distribution of kME values across gene sets, including the TIME_ACT signature genes.

### Single-sample gene-set enrichment (ssGSEA) analysis

TIME_ACT and comparator gene-expression signatures were quantified from uniformly normalized expression matrices using a gene-set-size–aware scoring strategy. To ensure consistency across heterogeneous transcriptomic and slide-inferred datasets, expression matrices were normalized independently within each cohort. Specifically, log-transformed TPM values (log₂(TPM + 1)) were converted to gene-wise z-scores across samples, thereby reducing gene-specific differences in baseline expression scale and technical variation while preserving relative expression differences between samples within each dataset. Genes with incomplete normalized values or very low variance were removed before signature scoring.

Because enrichment-based methods become unstable or uninformative when applied to very small gene sets, signature scoring was adapted according to signature size. Single-gene biomarkers were represented by the normalized z-score of the corresponding gene. Signatures containing 2-10 genes were scored as the arithmetic mean of the matched gene z-scores, which provides a direct and interpretable summary of coordinated expression when only a small number of genes are available. Any signatures containing more than 10 genes, including TIME_ACT, were quantified using single-sample gene-set enrichment analysis (ssGSEA). This threshold was used to ensure that ssGSEA was applied only to gene sets of sufficient size to support stable rank-based enrichment estimation, while avoiding the disproportionate influence of individual genes that can occur when enrichment methods are applied to very small signatures. ssGSEA was implemented using the GSVA package (v2.0.7) ^57^ in ssGSEA mode in R v4.4.3. The analysis was performed with alpha = 0, which assigns equal weight to all ranked positions, yielding an enrichment score that reflects the overall representation of signature genes across the expression rank, rather than emphasizing only the most highly expressed genes. The parameter normalize = TRUE was applied to rescale the scores to the range [–1, 1], enabling consistent comparisons across samples and tumor types.

### Spatial and Single-cell analysis of TIME_ACT signature genes

To investigate the spatial immune architecture, we analyzed two breast cancer tissue sections, formalin-fixed paraffin-embedded (FFPE) and fresh-frozen (FF), obtained from the 10x Genomics public dataset repository (https://www.10xgenomics.com/datasets). The FF tissue corresponded to HER2-positive, while the FFPE specimen displayed histological heterogeneity characterized by distinct regions of Ductal Carcinoma In Situ (DCIS) and invasive carcinoma of undetermined molecular subtype. Both tissue sections were accompanied by spot-level histopathological annotations provided by expert pathologists to categorize cellular components into tumor cells, stromal cells, lymphocytes, and macrophages. TIME_ACT scores were calculated for each individual spot, with spots exceeding the threshold of 0.0655 classified as “Hot” regions. Subsequently, spatial visualization maps were generated by overlaying pathologist-annotated cellular phenotypes with TIME_ACT-hot regions onto their corresponding tissue sections.

We obtained single-cell RNA-seq data from the GEO database (accession GSE115978), generated using the Smart-seq2 protocol on fresh tumor resections from 31 melanoma patients. The count matrix and accompanying metadata were imported into R and converted into a Seurat object. To compute cell-type– specific expression, we used Seurat’s “AverageExpression” function to calculate the mean log-TPM value of each gene within each annotated cell type. To mitigate inter-patient variability, we first calculated these cell-type–specific values for each patient individually and then averaged them across all patients for each gene and cell type. The resulting averaged expression matrix was visualized as a heatmap using the pheatmap package in R. Additionally, TIME_ACT gene expression patterns were retrieved from diverse single-cell RNA-seq datasets across multiple cancer types, as curated in the TISCH2 database ^58^.

### Predictive performance evaluation, threshold optimization, and statistical analysis

To evaluate the predictive performance of TIME_ACT, alongside comparator gene signatures and transcriptomic-based approaches, in stratifying responders from non-responders to immune checkpoint blockade (ICB) and other therapies, we computed the area under the receiver operating characteristic curve (AUC), area under the precision-recall curve (AUPRC) and the odds ratio (OR). AUC values were calculated using the roc_auc_score function, and AUPRC values were obtained using the average_precision_score function from the scikit-learn package (v1.4.2) ^59^ in Python 3.10, providing a threshold-independent measure of classification accuracy. A one-sided paired Wilcoxon signed-rank test was used to compare AUC values of TIME_ACT against each signature and method across cohorts. To define a consistent classification threshold for pan-cancer TIME_ACT stratification, we applied Youden’s index ^60^ (J = Sensitivity + Specificity – 1) across nine TCGA cancer types using predefined Hot and Cold tumor labels. The TIME_ACT score that maximized Youden’s index was selected as the optimal pan-cancer decision threshold (0.0655). This threshold was then applied uniformly across all external validation cohorts to classify samples as TIME_ACT-high or TIME_ACT-low and to compute odds ratios for treatment response. All additional statistical analyses, including Pearson correlation and multiple hypothesis testing (Benjamini–Hochberg false discovery rate), were performed in R v4.4.3.

### Clinical Histopathology Cohorts

To evaluate the TIME_ACT signature in real-world clinical settings, we curated a multi-institutional collection of new and unpublished clinical histopathology datasets encompassing whole-slide H&E-stained tumor sections alongside detailed treatment annotations and clinical outcomes. In total, we collected eight cohorts from seven cancer types, each with annotated treatment information and clinical response to therapy. One additional dataset was received that did not include treatment response information; only overall survival data were available for the patients. We received slides and metadata with variable levels of detail and formatting. Across all datasets, we only retained slides from patients who received ICB as part of their treatment regimen. Here is the description of each cohort that we analyzed.

1. ***NCI Bladder Cancer (urothelial carcinoma) cohort:*** The NCI bladder cancer (BLCA) cohort included 48 patients (48 slides) with locally advanced or metastatic urothelial carcinoma of the bladder. This was a subset of a larger trial (NCT02496208) that enrolled 150 patients across multiple metastatic genitourinary tumors (GU). All the BLCA patients included in our analysis were treated with cabozantinib and nivolumab alone or with ipilimumab. All slides were archival pre-treatment FFPE specimens. In total, 17 patients responded to therapy and 31 did not.
2. ***Cedars-Sinai Melanoma (SKCM) cohort:*** This cohort comprised 32 patients with advanced melanoma (32 slides). All patients were treated with ICB regimens. Treatments included anti-PD-1 monotherapy in some cases and various combination immunotherapies - for example, PD-1 inhibitors combined with CTLA-4 blockade (ipilimumab + nivolumab), with LAG-3 blockade, or with other investigational immunomodulators (such as an IDO1 inhibitor or GITR agonist) - reflecting the clinical trial therapies used. There were 18 responders and 14 non-responders in this melanoma cohort.
3. ***Cedars-Sinai Breast Cancer (BRCA) cohort***: This cohort included a total of 23 tumor slides, derived from the Cedars-Sinai “Molecular Twin Research Umbrella Protocol” (IRB STUDY00001879). All patients had advanced invasive breast carcinoma and were treated with ICB (anti-PD-1/PD-L1 therapy). Of the 23 samples, 15 were annotated as responders and 8 as non-responders. In the responder group, 11 were treated with a combination of ICB and chemotherapy, and four patients received a combination of immunotherapy (pembrolizumab and trastuzumab) and supportive chemotherapy. In the non-responder group, 6 received ICB and chemotherapy (in one patient, chemotherapy, ICB, and trastuzumab), and two received ICB therapy alone. H&E slides were derived from surgical FFPE specimens collected between January 2013 and July 2024.
4. ***Cedars-Sinai Lung Adenocarcinoma (LUAD) cohort:*** The Cedars-Sinai LUAD cohort consisted of 30 patients (30 slides in total), all with KRAS-mutant lung adenocarcinoma. All patients had stage IV or recurrent lung adenocarcinoma and received ICB, either alone or in combination with chemotherapy. The majority (24/30) were treated with first-line platinum-based chemotherapy plus anti-PD-1 immunotherapy, while 6 patients received immunotherapy alone. 14 patients were responders, and 16 patients were non-responders. All slides were derived from surgical FFPE specimens.
5. ***HCRN Renal Cell Carcinoma cohort:*** This cohort included 96 patients with metastatic clear cell renal cell carcinoma (KIRC; 151 pre-treatment tumor slides) and 28 patients with advanced papillary renal cell carcinoma (KIRP; 48 pre-treatment tumor slides). All patients were enrolled in the HCRN GU16-260 clinical trial (NCT03117309) and received nivolumab monotherapy. Multiple patients contributed paired specimens from primary and metastatic sites. Among the KIRC patients, 35 responded to therapy and 61 did not. Among the KIRP patients, 4 were responders and 24 were non-responders. All slides were derived from FFPE tissue blocks. A subset of KIRC de-identified RNA-seq data, linked to histopathology and clinical outcome, was retrieved through dbGaP (phs003618.v1.p1).
6. ***Taiwan-NYCU Head and Neck Squamous Cell Carcinoma (HNSCC) cohort:*** The Taiwan-NYCU dataset was collected under the project of the National Science and Technology Council, Taiwan, and includes head and neck squamous cell carcinoma (HNSCC) patients treated at Taipei Veterans General Hospital. The dataset was acquired from the Big Data Center of Taipei Veterans General Hospital. Diagnostic slides were derived from surgical FFPE specimens collected as part of standard clinical care between July 2021 and July 2024. Cohort 1 includes 33 HNSCC patient slides, among whom 11 were classified as responders to immunotherapy and 22 as non-responders. Among responders, 8 received ICB monotherapy, 2 received ICB with chemotherapy, and 1 received ICB with targeted therapy. Among non-responders, 5 received ICB monotherapy, 11 received ICB with chemotherapy, 5 received ICB with targeted therapy, and 1 received the full combination. Cohort 2 includes another 62 HNSCC patient slides. Within this cohort, 21 patients were identified as responders, and 41 as non-responders. Among responders, 5 received ICB monotherapy, 8 received ICB with chemotherapy, 2 received ICB with targeted therapy, and 6 received the full combination. In the non-responder group, 19 received ICB monotherapy, 14 received ICB with chemotherapy, 5 received ICB with targeted therapy, and 3 received the full combination. We used 61 slides, not included one non-responder patient slide due to poor image quality.
7. ***Taiwan-NYCU Liver Cancer (LIHC) cohort:*** This cohort included 30 patients with hepatocellular carcinoma treated with ICB at Taipei Veterans General Hospital between 2021 and 2024. Among them, 10 patients were responders, and 20 were non-responders. All analyses were performed on pre-treatment H&E-stained FFPE slides.

### Inferring gene expression from histopathology images using Path2Omics

We utilized our recently developed Path2Omics pipeline ^36^ to infer gene expression profiles directly from pathology slide images. Briefly, Path2Omics consists of three main components: image pre-processing, feature extraction, and regression. In the image pre-processing step, we first divided each whole slide image into non-overlapping tiles of 512 x 512 RGB pixels. We then applied Sobel edge detection to select tiles containing tissue ^61^. To minimize staining variation (heterogeneity and batch effects) between institutions, we applied Macenko’s method for color normalization to the selected tiles ^62^. For feature extraction, a foundational model pre-trained on digital pathology data was used to derive informative features from each tile. These features were then input into a multi-layer perceptron within the regression module to predict gene expression values. Path2Omics was initially trained and validated on the TCGA dataset using 5-fold cross-validation, separately for each cancer type. The pre-trained models were subsequently applied to predict gene expression from slides on the clinical treatment datasets, without additional training. The methodological details are provided in the original manuscript ^36^.

### Model for predicting treatment response directly from slides

The “direct” model was developed to predict patient response directly from the slide, without the intermediate step of inferring gene expression and TIME_ACT score. This model utilized the same image pre-processing and feature extraction framework as Path2Omics but replaced the regression component with a classification module. Since the response labels were only available at the slide level, a weakly supervised training scheme was implemented, where all tiles from a slide were assigned the corresponding slide label. Slide-level predictions were then derived by averaging the tile-level outputs. In contrast to the “indirect” approach, where Path2Omics, pre-trained on TCGA data, was used to predict gene expression in the treatment cohort, without further fine-tuning, the “direct” model was fully trained and validated on the same treatment cohort using a 5-fold cross-validation, as it required response labels for supervision.

### Data availability

TCGA data are publicly available at the Genomic Data Commons Data Portal (https://portal.gdc.cancer.gov). Details of the 25 publicly available ICB transcriptomic datasets are provided in Supplementary Table 6. Requests for access to unpublished data from nine clinical histopathology cohorts should be directed to the corresponding authors. Each request will be assessed individually within 15 business days to ensure adherence to intellectual property rights and patient privacy standards. All TIME_ACT scores and corresponding response information for each sample for both transcriptomic and histopathology cohorts are available in the Zenodo repository (https://zenodo.org/records/22313248).

### Code availability

The code supporting this publication will be made available for academic research purposes via: (https://zenodo.org/records/22313248). Any additional information is available from the corresponding authors upon request.

## Acknowledgements

This research is supported in part by the Intramural Research Program of the National Institutes of Health, National Cancer Institute, Center for Cancer Research. This work has utilized the computational resources of the NIH HPC Biowulf cluster (http://hpc.nih.gov). We thank the Cedars-Sinai Medical Center OncoBiobank and Pathology Shared Resources for their technical assistance. We also thank the Big Data Center of Taipei Veterans General Hospital for their help in data sharing. The results presented are in part based on the data generated by the Cancer Genome Atlas Research Network (https://www.cancer.gov/tcga/). This research is also partially supported by a grant of the Korea-US Collaborative Research Fund (KUCRF), funded by Ministry of Science and ICT and Ministry of Health & Welfare, Republic of Korea (grant number: RS-2024-00468417). We would additionally like to acknowledge Nicholas P. Restifo, Nishanth Ulhas Nair, Sanna Madan, and Alejandro Schaffer for helpful discussions, and the members of the Cancer Data Science Laboratory for their constructive feedback. ChatGPT v5.3 (https://chatgpt.com/) was used strictly to refine certain existing paragraphs to improve sentence constructions.

## Supplementary Tables Descriptions

**Supplementary Table 1 |** Differentially expressed genes between immune-hot and immune-cold TCGA melanoma samples. This table lists the 339 genes significantly upregulated in TIME_ACT-defined immune-hot tumors versus cold tumors in the TCGA-SKCM cohort, identified using limma with stringent thresholds on both log₂ fold change (logFC > 3.344) and B-statistic (B > 36.224). Includes gene identifiers, average expression, t-statistics, p-values, adjusted p-values, and B-statistics.

**Supplementary Table 2 |** Gene ontology enrichment analysis of genes upregulated in immune-hot tumors. Enriched immune-related biological processes among the 339 upregulated genes in TIME_ACT-hot tumors. Columns include GO term name, adjusted p-value, odds ratio, and combined enrichment score from Enrichr.

**Supplementary Table 3 |** The TIME_ACT 66-gene immune activation signature. List of 66 genes forming the TIME_ACT signature, derived from genes upregulated in immune-hot tumors and involved in top-ranked immune-related GO terms.

**Supplementary Table 4 |** TIME_ACT scores and immune activation annotations across annotated TCGA cohorts. TIME_ACT scores and their associated Hot/Cold immune annotations for TCGA samples from the 9 tumor types. Includes sample ID, TIME_ACT score, annotation, and cancer type.

**Supplementary Table 5 |** TIME_ACT scores across all TCGA tumors. Pan-cancer TIME_ACT scores computed for all TCGA tumor samples using ssGSEA. This table provides sample ID, TIME_ACT score, and tumor type.

**Supplementary Table 6 |** Summary of pre-treatment anti-PD1 transcriptomic cohorts used in this study. Overview of the 25 ICB-treated transcriptomic datasets used to assess the predictive performance of TIME_ACT and to compare it with other signatures and methods. Includes cohort name, cancer type, total sample size, number of responders, and non-responders.

**Supplementary Table 7 |** List of comparator immune-related gene expression signatures used for benchmarking. Details of 21 established immune-related transcriptomic signatures used for benchmarking TIME_ACT. Includes signature name, type, PubMed ID (PMID), and gene list for each signature.

---

**Supplementary Fig. 1.**
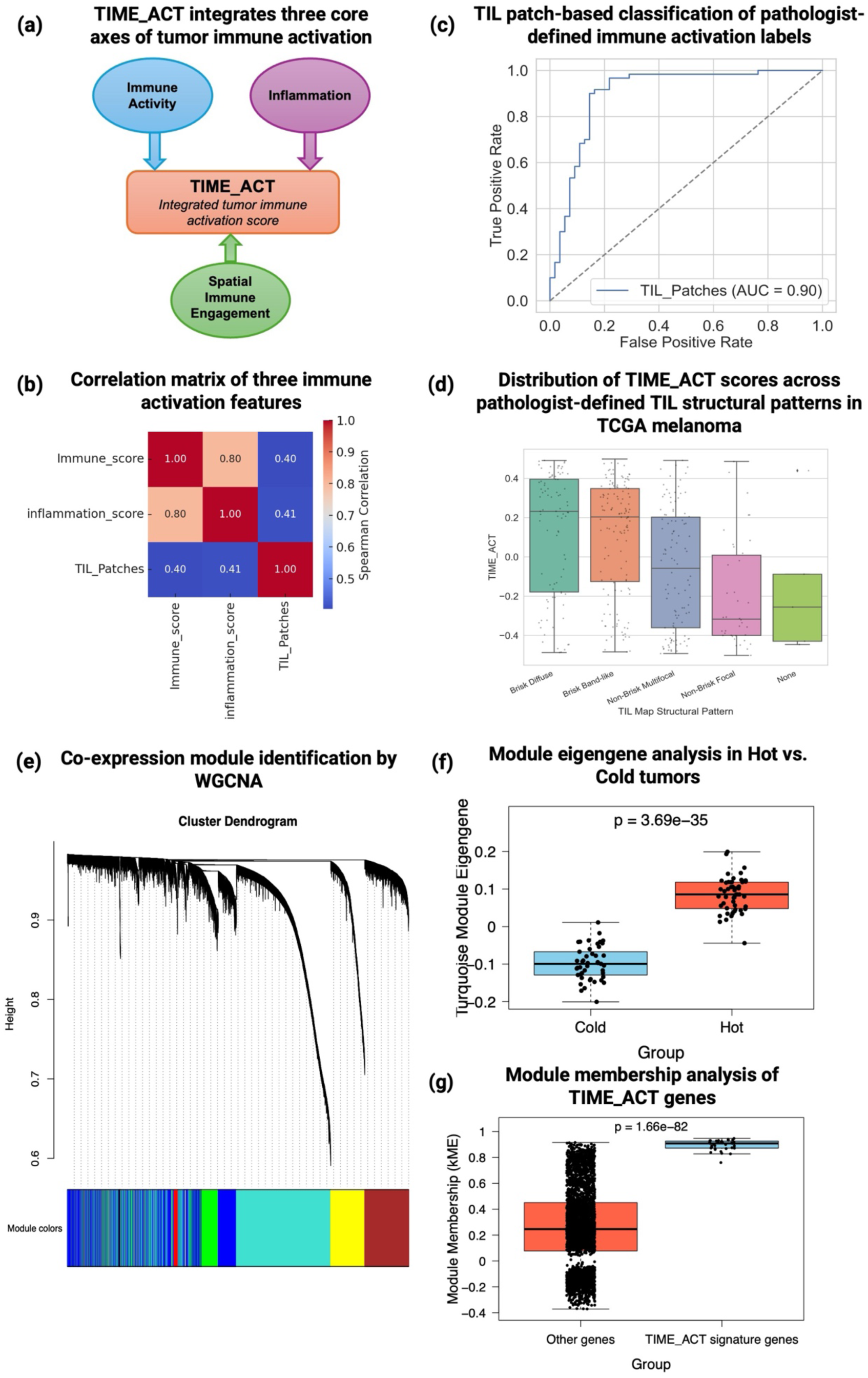
TIME_ACT integrates multi-dimensional immune activation and defines a co-expressed immune module in melanoma. **(a)** Schematic illustrating the conceptual design of TIME_ACT, which integrates three major immune activation dimensions, inflammation, immune activity, and spatial immune engagement into a unified tumor immune activation score. **(b)** Spearman correlation matrix showing positive associations among the three axes of immune activation across TCGA melanoma samples. **(c)** ROC curve showing the performance of TIL patch counts (derived from Saltz et al.) in discriminating pathologist-defined hot versus cold TCGA-SKCM tumors. The TIL patch metric achieves a high classification accuracy (AUC = 0.90), indicating strong concordance between histopathology image-derived lymphocyte spatial features and expert histopathologic assessment of tumor immune activation. **(d)** Boxplots show the distribution of TIME_ACT scores across the five pathologist-defined TIL Map structural patterns in TCGA-SKCM, as defined by Saltz et al. Each point represents an individual tumor sample. TIME_ACT scores are highest in tumors with brisk lymphocytic infiltration and progressively decrease across non-brisk patterns, reaching the lowest levels in TIL-absent tumors. This trend demonstrates that the transcriptomic TIME_ACT signature closely reflects the degree and structural organization of lymphocyte infiltration observed in routine H&E histopathology. **(e)** Hierarchical clustering dendrogram of genes from TCGA melanoma tumors grouped into distinct co-expression modules by WGCNA. Each color in the horizontal bar represents a different gene module. The turquoise module, highlighted in the plot, contains all 66 genes from the TIME_ACT signature, indicating that they are tightly co-expressed within a single immune-related network module. **(f)** Boxplot comparing the eigengene values of the turquoise module between cold and hot tumors. The turquoise module shows significantly higher expression in hot tumors, indicating a strong association with immunologically active TIME. **(g)** Module membership (kME) scores comparing TIME_ACT signature genes vs. all other genes within the turquoise module. TIME_ACT genes exhibit significantly stronger module membership, highlighting their central role in the hot immune phenotype co-expression network.

**Supplementary Fig. 2.**
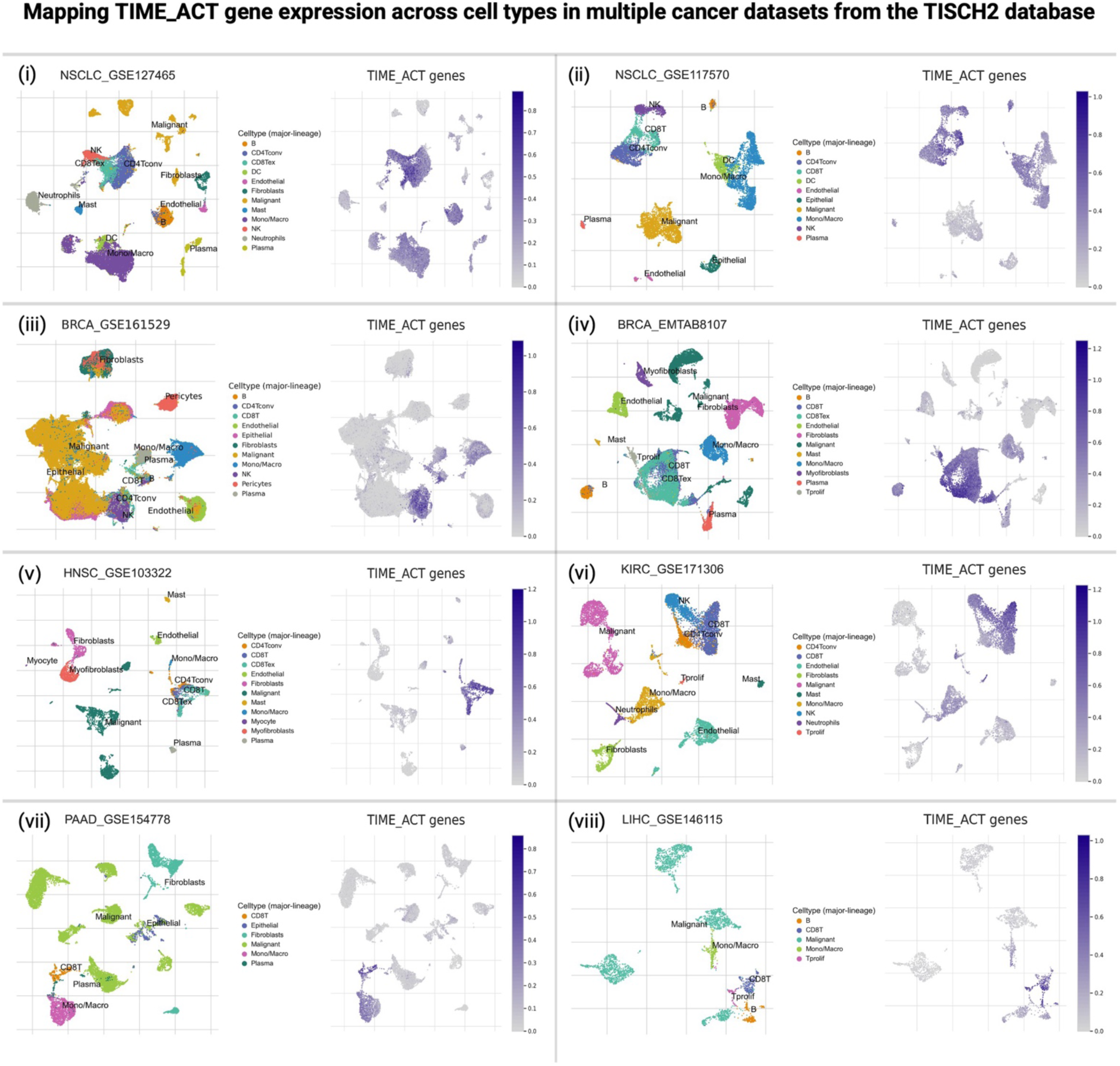
TIME_ACT genes are consistently expressed in immune cell populations across multiple cancer types. Cell-type-specific expression of TIME_ACT genes was assessed using eight annotated single-cell RNA-seq datasets from the TISCH2 database. (i-viii) UMAP plots and gene expression heatmaps demonstrate consistent enrichment of TIME_ACT gene expression in immune compartments, particularly CD8⁺ T cells, CD4⁺ T cells, and B cells, across lung cancer, breast cancer, liver cancer, and other tumor types. Minimal or no expression was observed in malignant cells, fibroblasts, or endothelial cells. These findings validate the immune cell specificity of the TIME_ACT signature and reinforce its relevance as a marker of functional immune engagement across diverse TME.

**Supplementary Fig. 3.**
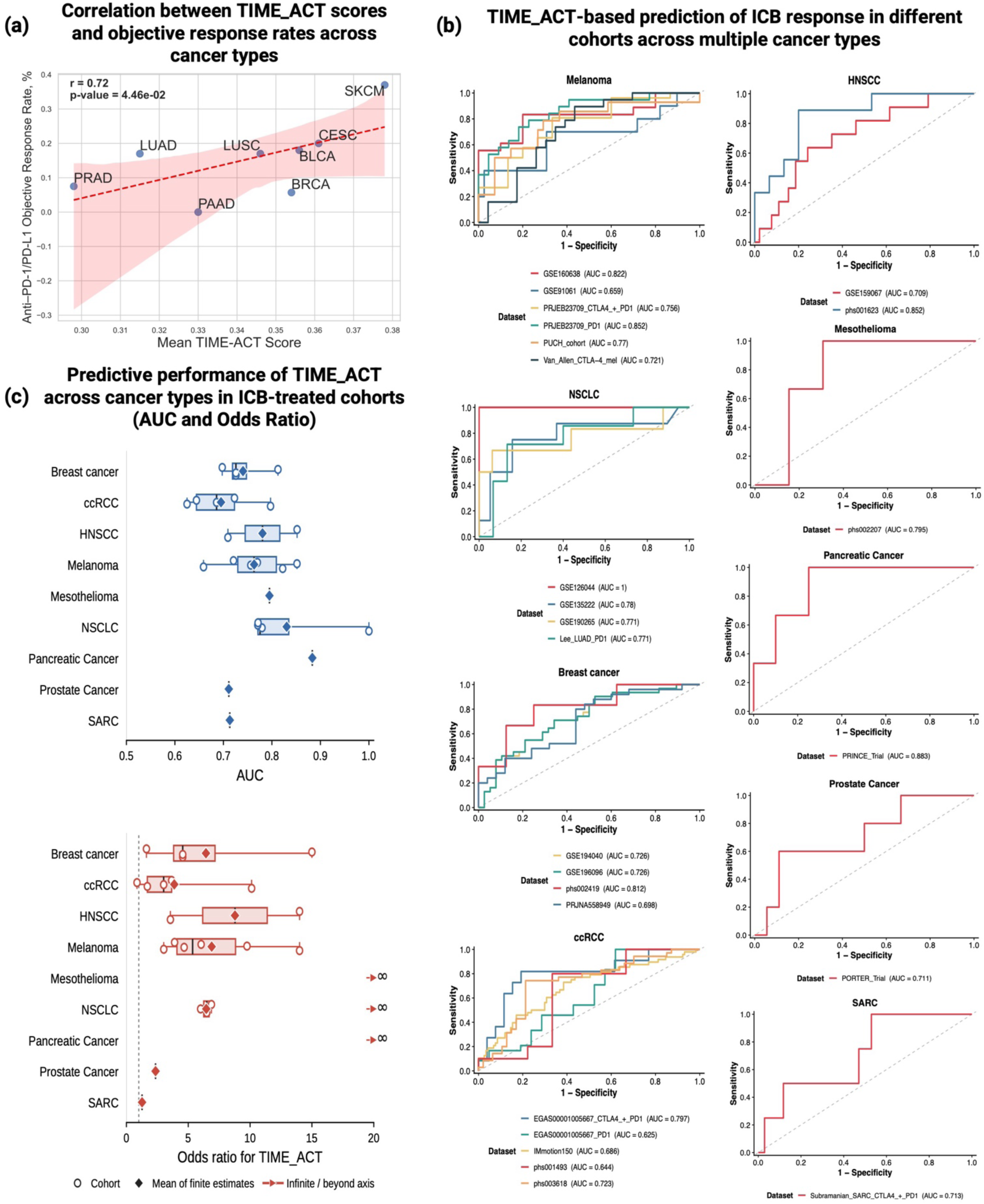
Pan-cancer association and predictive performance of TIME_ACT in ICB-treated cohorts. **(a)** Correlation between mean TIME_ACT scores in Hot tumors and objective response rates (ORRs) to ICB therapy across eight cancer types with available data. Higher immune activation, as captured by TIME_ACT, is associated with higher ORR. **(b)** Receiver operating characteristic (ROC) curves showing the performance of pretreatment TIME_ACT scores for predicting ICB response across 25 independent cohorts spanning nine cancer types. Curves are grouped by cancer type, and cohort-specific AUC values are indicated in the legends. **(c)** Cohort-level predictive performance of TIME_ACT across 25 independent immune checkpoint blockade (ICB) pre-treatment transcriptomic cohorts. AUC (top) and odds ratio (OR; bottom) are shown for each cohort across cancer types. Points denote individual cohorts, box plots summarize cohort distributions, and OR values exceeding the plotting range are indicated by ∞ at the axis boundary. TIME_ACT exhibited consistently strong discrimination and enrichment of responders across multiple cancer types, including melanoma, NSCLC, HNSCC, and breast cancer.

**Supplementary Fig. 4.**
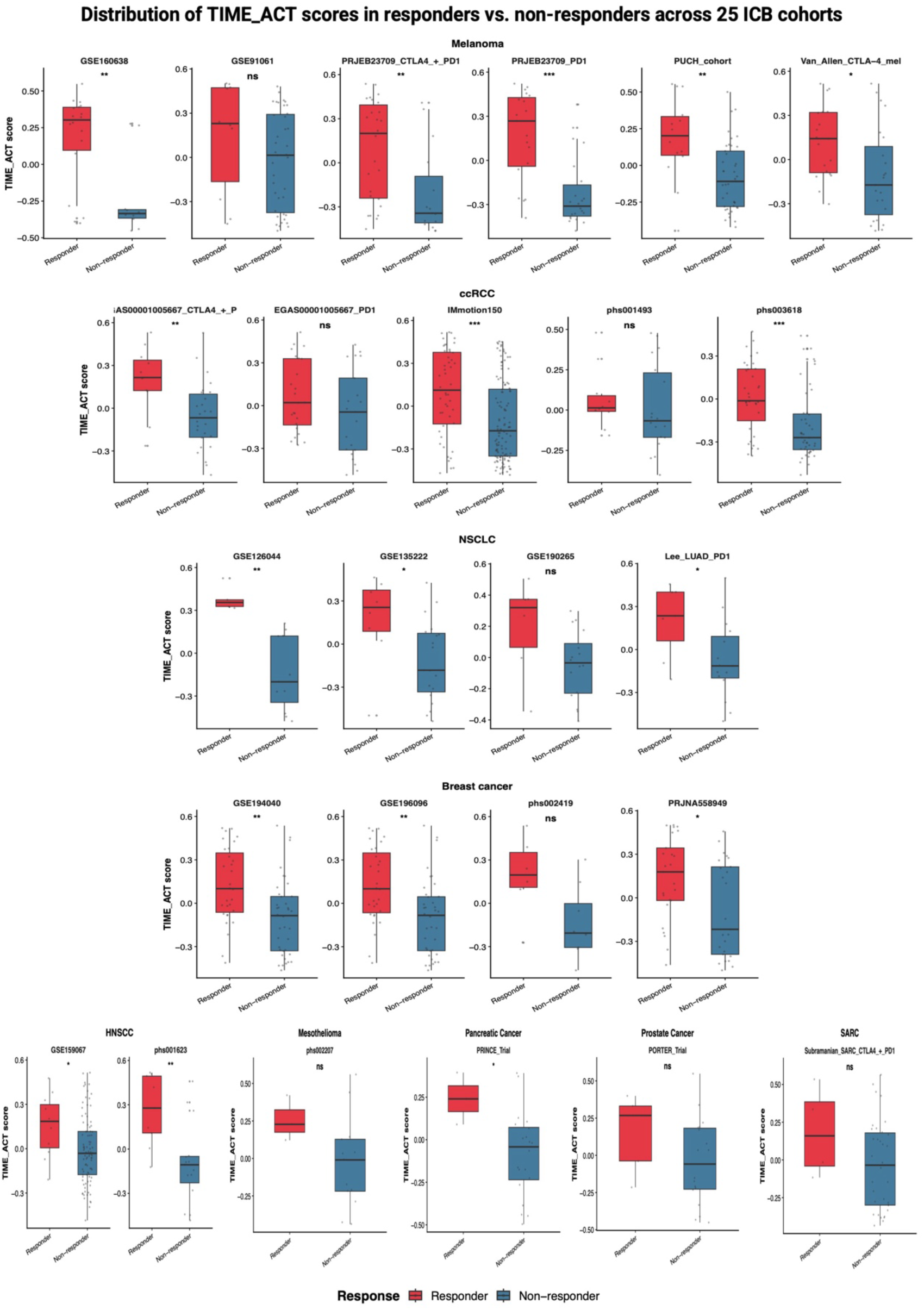
Distribution of TIME_ACT scores between responders and non-responders across independent ICB-treated cohorts. Box plots show TIME_ACT scores in responders and non-responders across 25 independent ICB cohorts. Each point represents an individual patient. Responders consistently display significantly higher TIME_ACT scores, underscoring the predictive value of the signature in clinically relevant settings. Statistical significance was assessed within each cohort using a two-sided Wilcoxon rank-sum test. *P < 0.05 (*), P < 0.01 (**)* and *P < 0.001 (***)*; *ns*, not significant.

**Supplementary Fig. 5.**
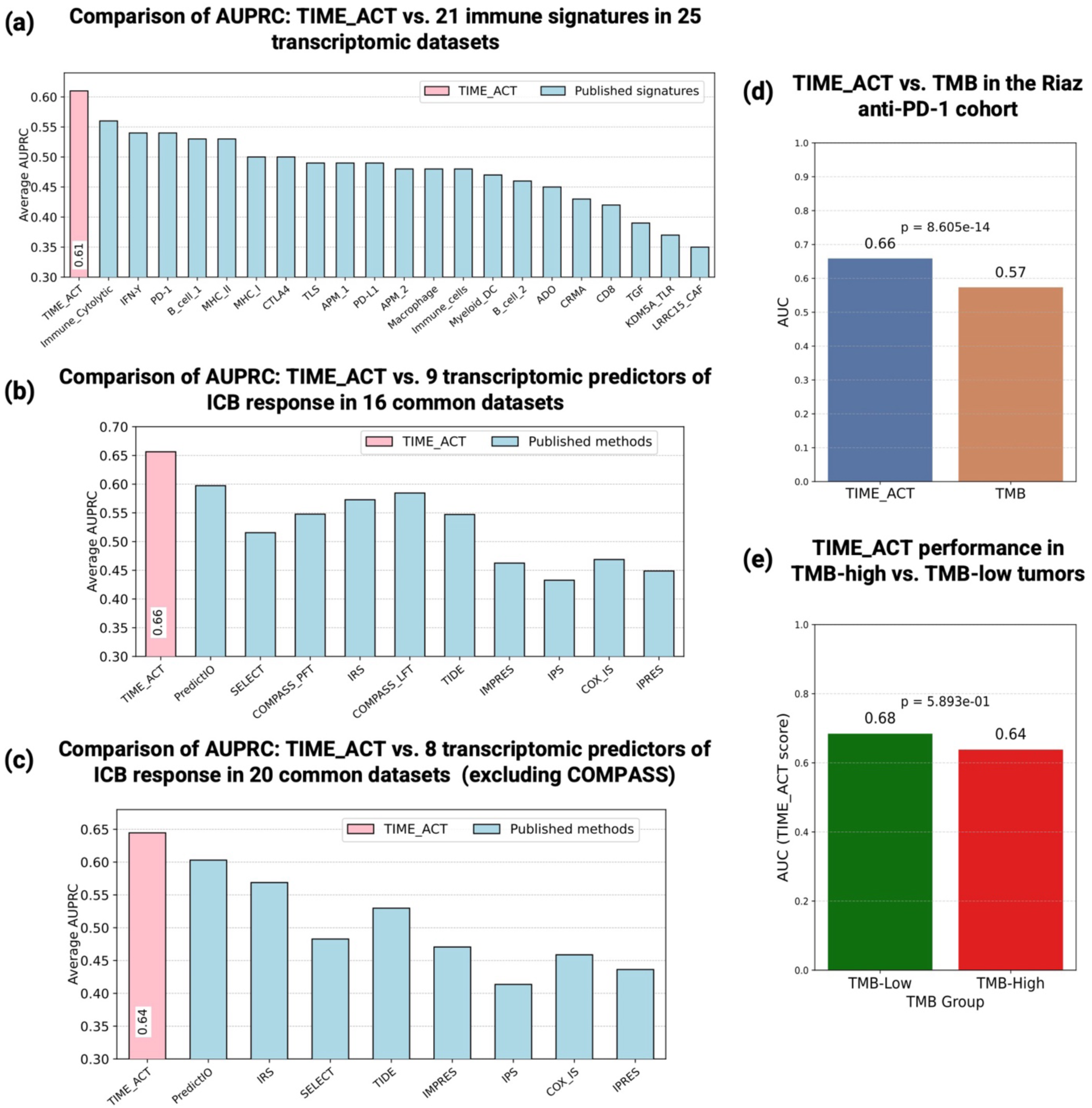
Benchmarking TIME_ACT against immune-related signatures, transcriptomic methods using precision-recall performance, and also with TMB in predicting ICB response. (a) Comparison of the mean area under the precision-recall curve (AUPRC) for TIME_ACT and 21 published immune-related gene-expression signatures across 25 independent pretreatment ICB cohorts. TIME_ACT achieved the highest mean AUPRC. (b) Head-to-head comparison of TIME_ACT with nine transcriptomic ICB response prediction methods across the 16 cohorts in which all methods, including COMPASS-PFT and COMPASS-LFT, could be evaluated. Bars indicate the mean AUPRC across cohorts. (c) Comparison of TIME_ACT with eight transcriptomic prediction methods across 20 common cohorts after excluding COMPASS, which could not be reliably applied to the additional cohorts because of insufficient coverage of its required input genes. TIME_ACT retained the highest mean AUPRC in both matched comparisons. (d) In the Riaz et al. melanoma cohort with matched RNA-seq and TMB data, TIME_ACT (AUC = 0.66) outperforms TMB (AUC = 0.57) in predicting ICB response. (e) TIME_ACT retains predictive power in both TMB-High (AUC = 0.64) and TMB-Low (AUC = 0.68) tumors, demonstrating its utility in genomic biomarker-poor settings where TMB fails to stratify responders.

**Supplementary Fig. 6.**
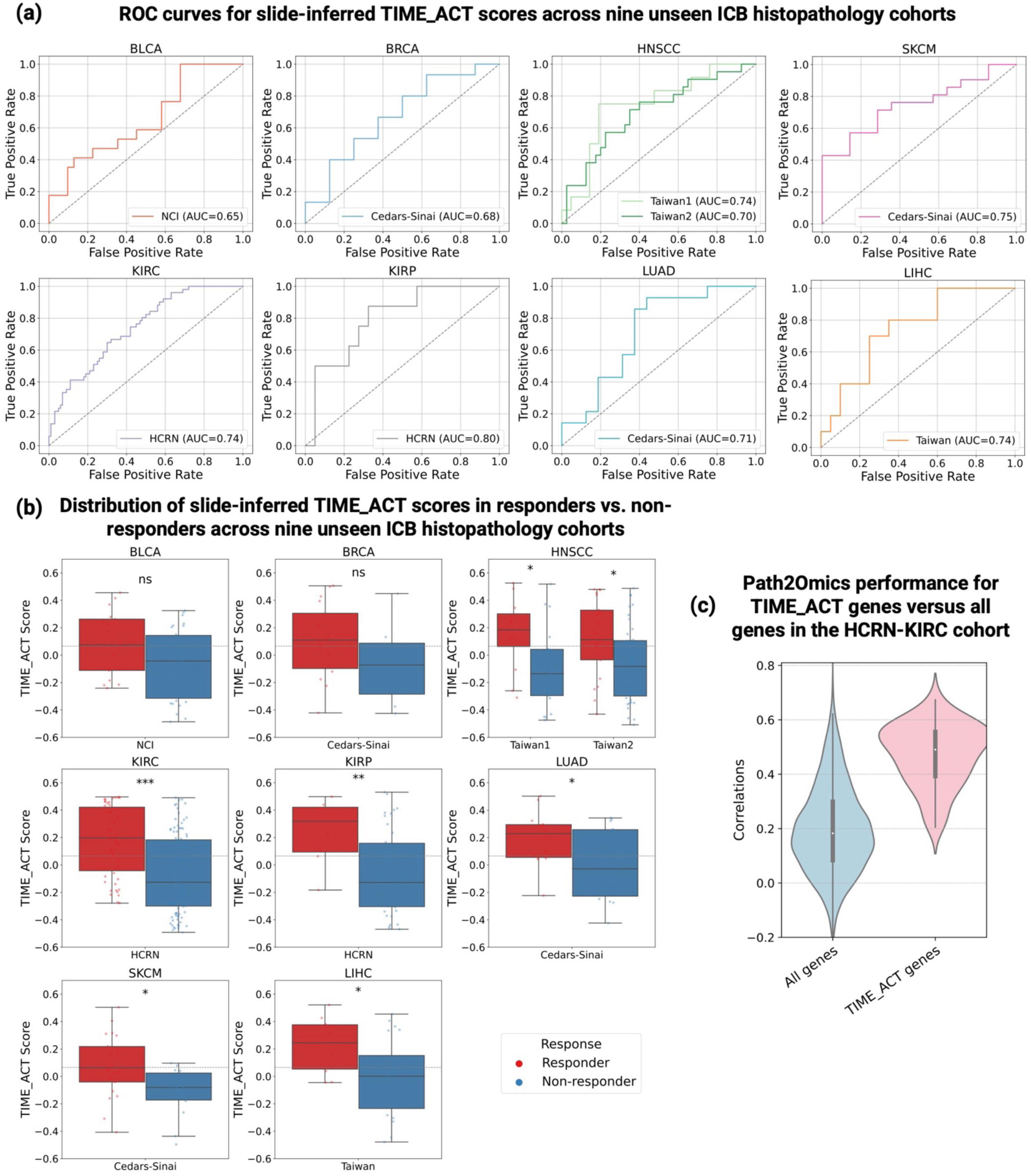
TIME_ACT scores inferred from histopathology predict ICB response across nine unseen cohorts. **(a)** ROC curves evaluating the predictive performance of slide-inferred TIME_ACT scores in nine independent ICB-treated cohorts. TIME_ACT scores were computed using gene expression inferred from diagnostic H&E slides via the Path2Omics model. AUC values ranged from 0.65 to 0.80, with a mean AUC of 0.73, indicating consistent predictive performance across cancer types and institutions. **(b)** Box plots showing the distribution of slide-inferred TIME_ACT scores in responders versus non-responders. In all eight cohorts, responders exhibit significantly higher inferred TIME_ACT scores, reinforcing the translational relevance of histology-based immune activation prediction for clinical decision-making. **(c)** Path2Omics performance for TIME_ACT genes versus all genes in the HCRN-KIRC cohort. Violin plots demonstrate the distribution of Pearson correlation coefficients between slide-inferred and measured RNA-seq expression for TIME_ACT genes (pink) versus all genes (light blue) across samples. Notably, these results were obtained by using the Path2Omics model pre-trained on TCGA and applied directly without any fine-tuning.

