## Supplementary figures and images for "Pan-cancer prediction of tumor immune activation and response to immune checkpoint blockade from tumor transcriptomics and histopathology"

### Supplementary Fig.1

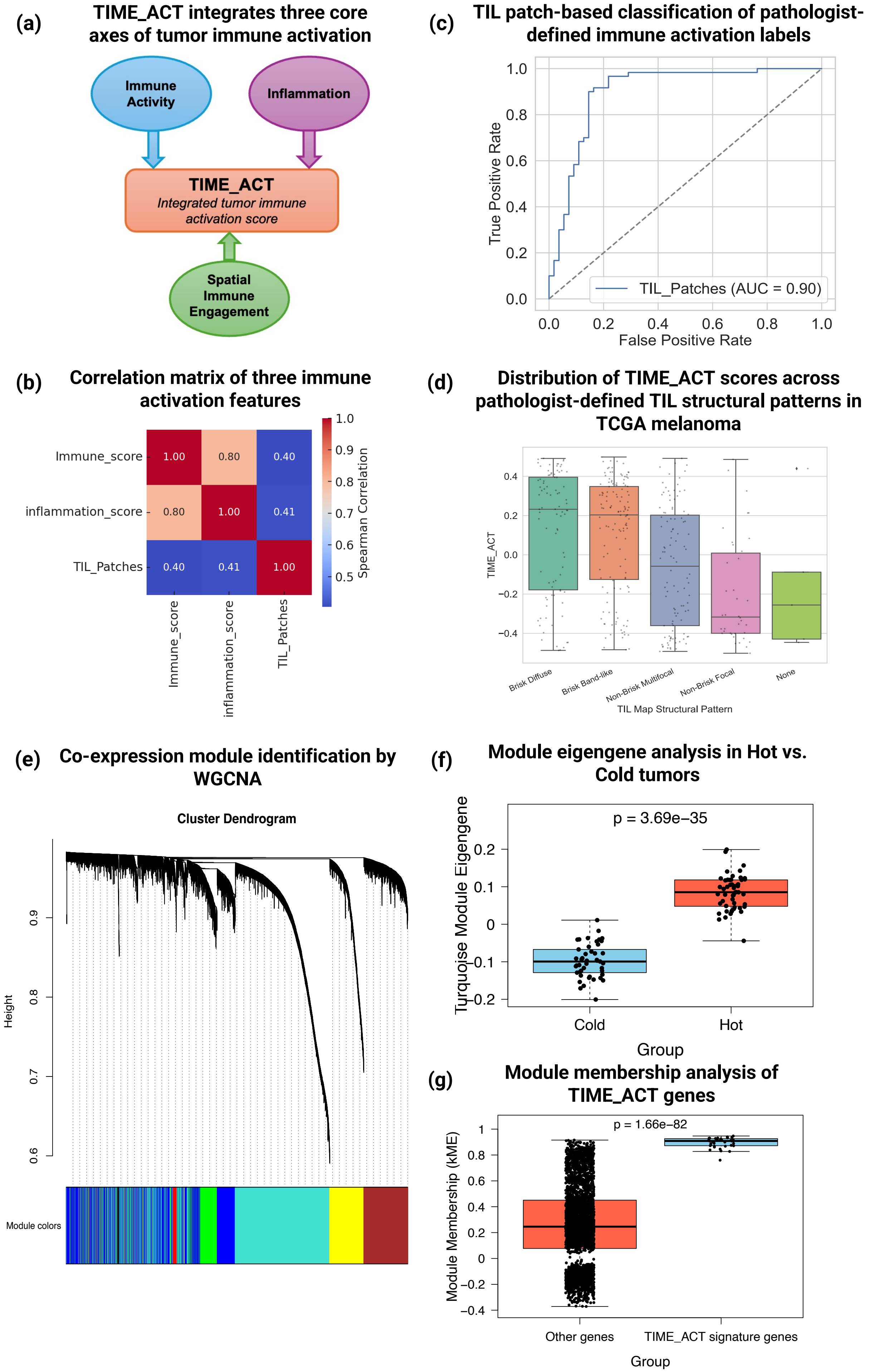

### Supplementary Fig.3

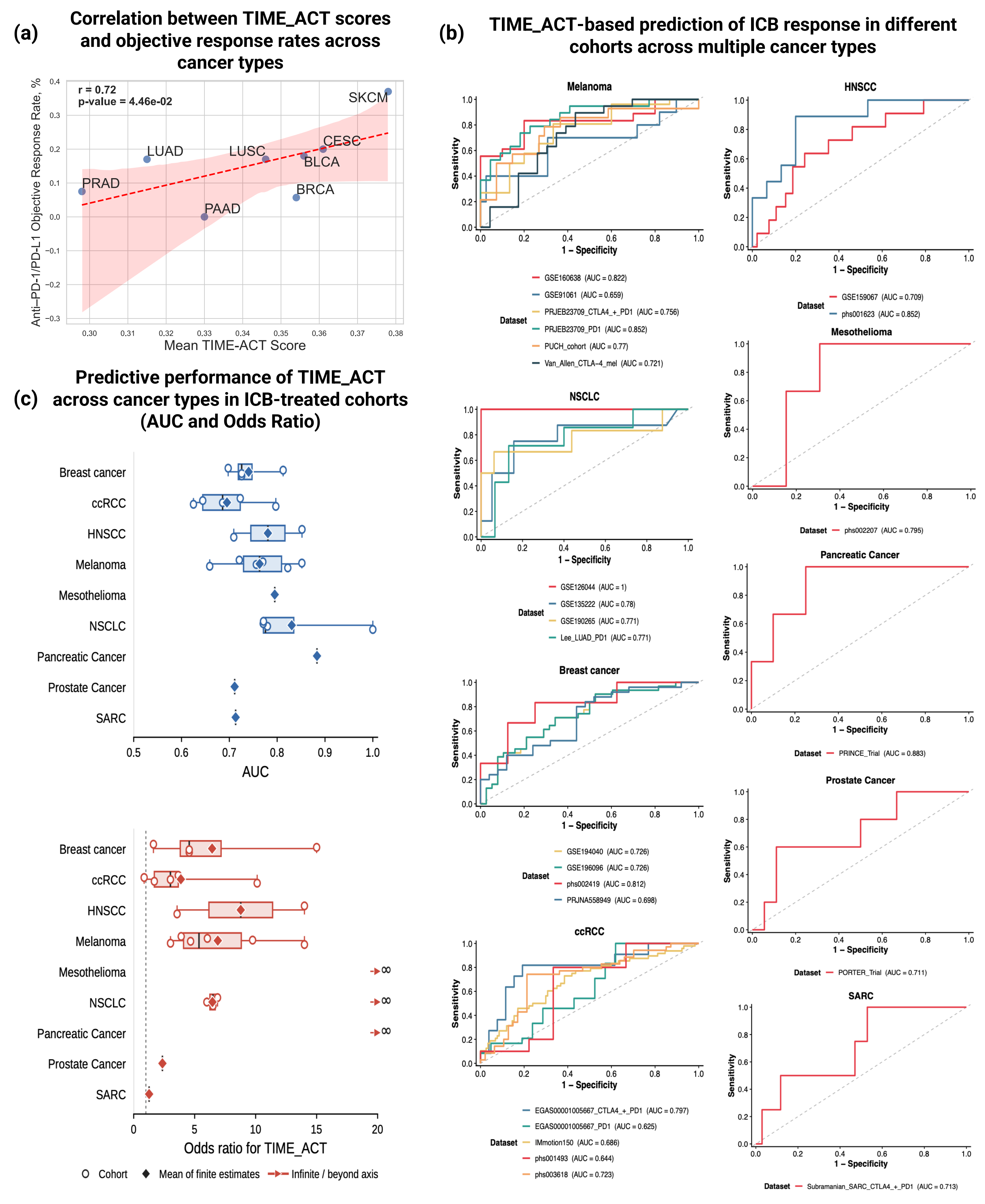

### Supplementary Fig.4

#### Distribution of TIME_ACT scores in responders vs. non-responders across 25 ICB cohorts

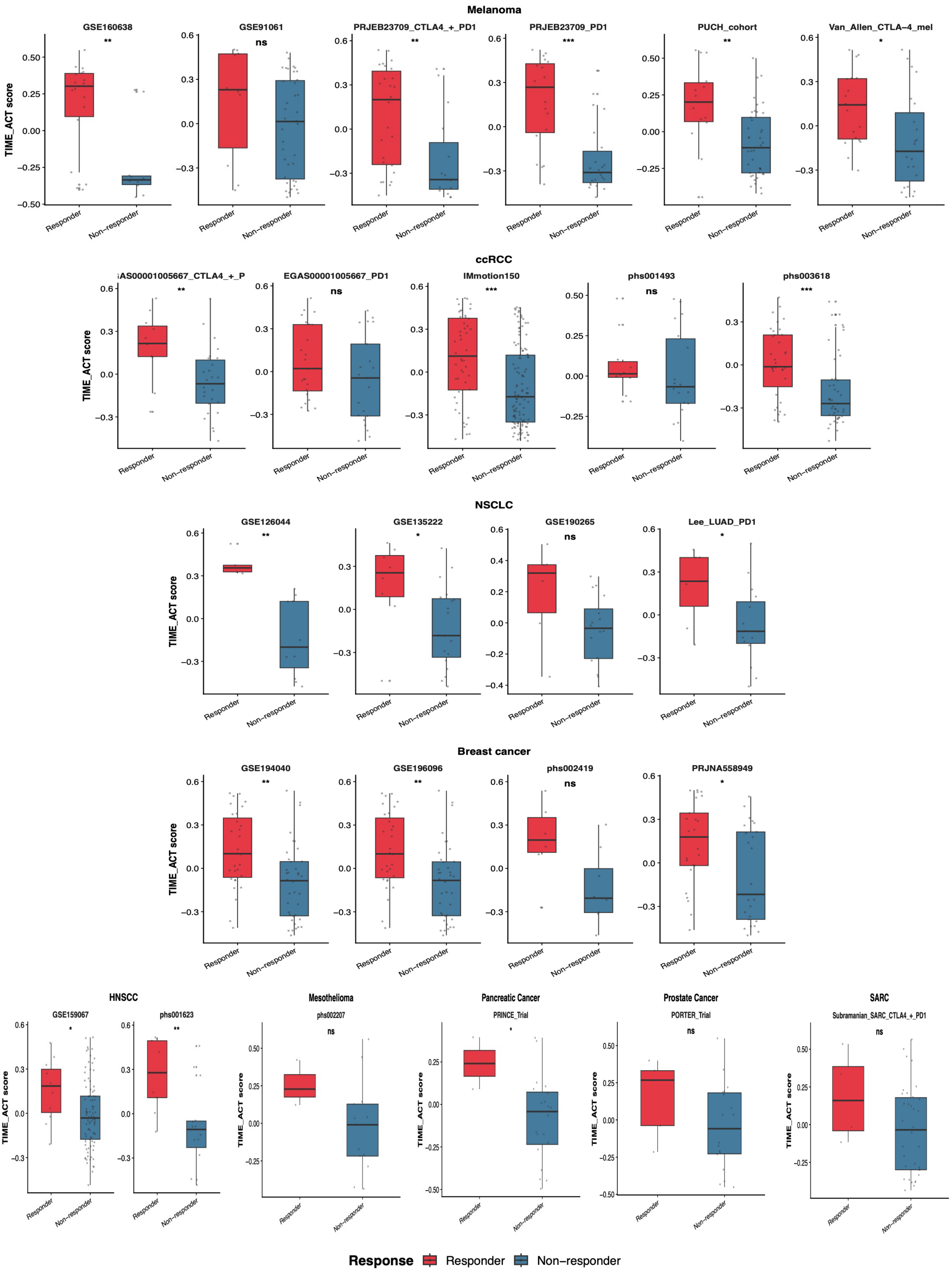

### Supplementary Fig.5

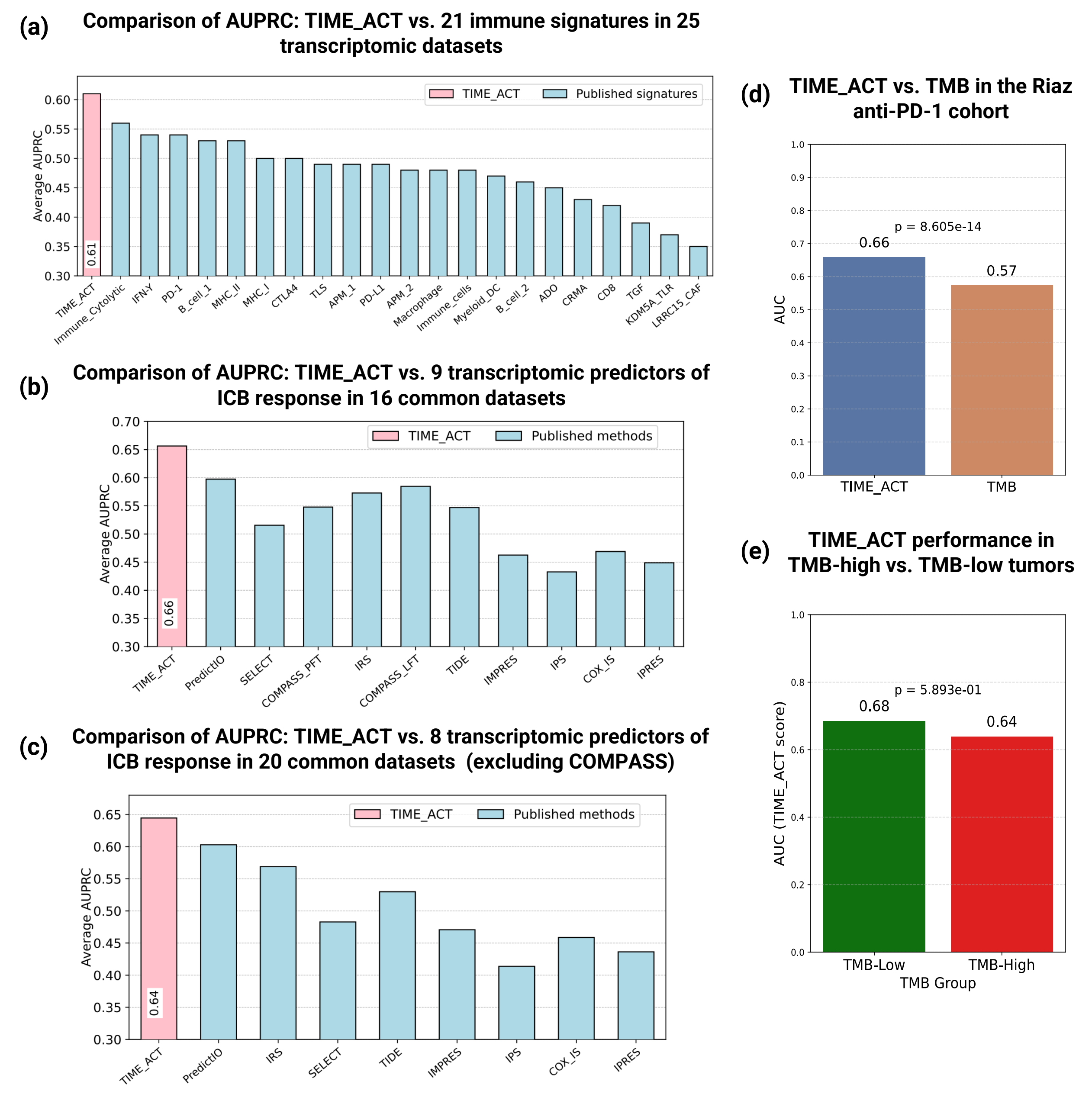

### Supplementary Fig.6

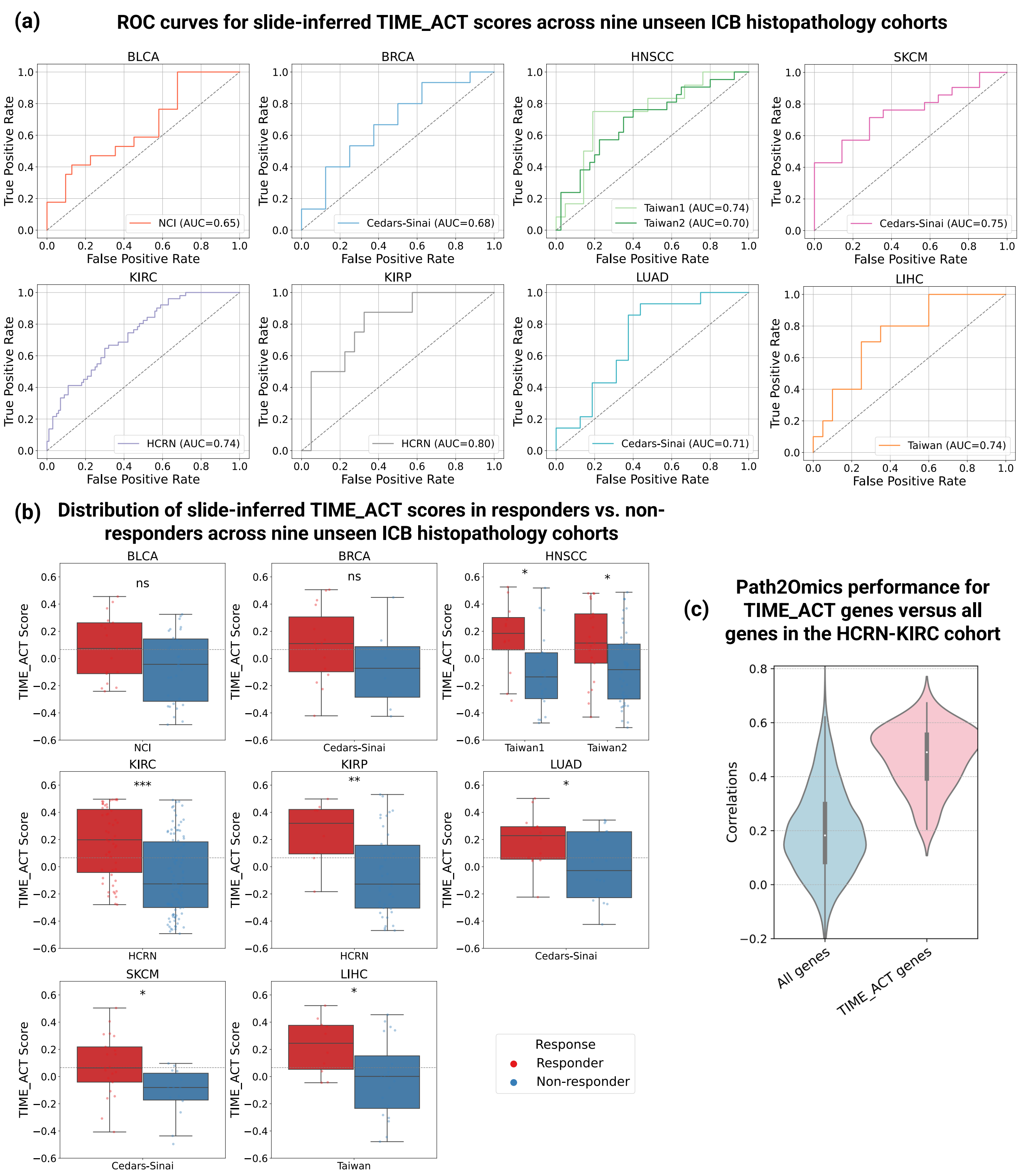
