## Supplementary Fig.2 for "Pan-cancer prediction of tumor immune activation and response to immune checkpoint blockade from tumor transcriptomics and histopathology"

### Mapping TIME_ACT gene expression across cell types in multiple cancer datasets from the TISCH2 database

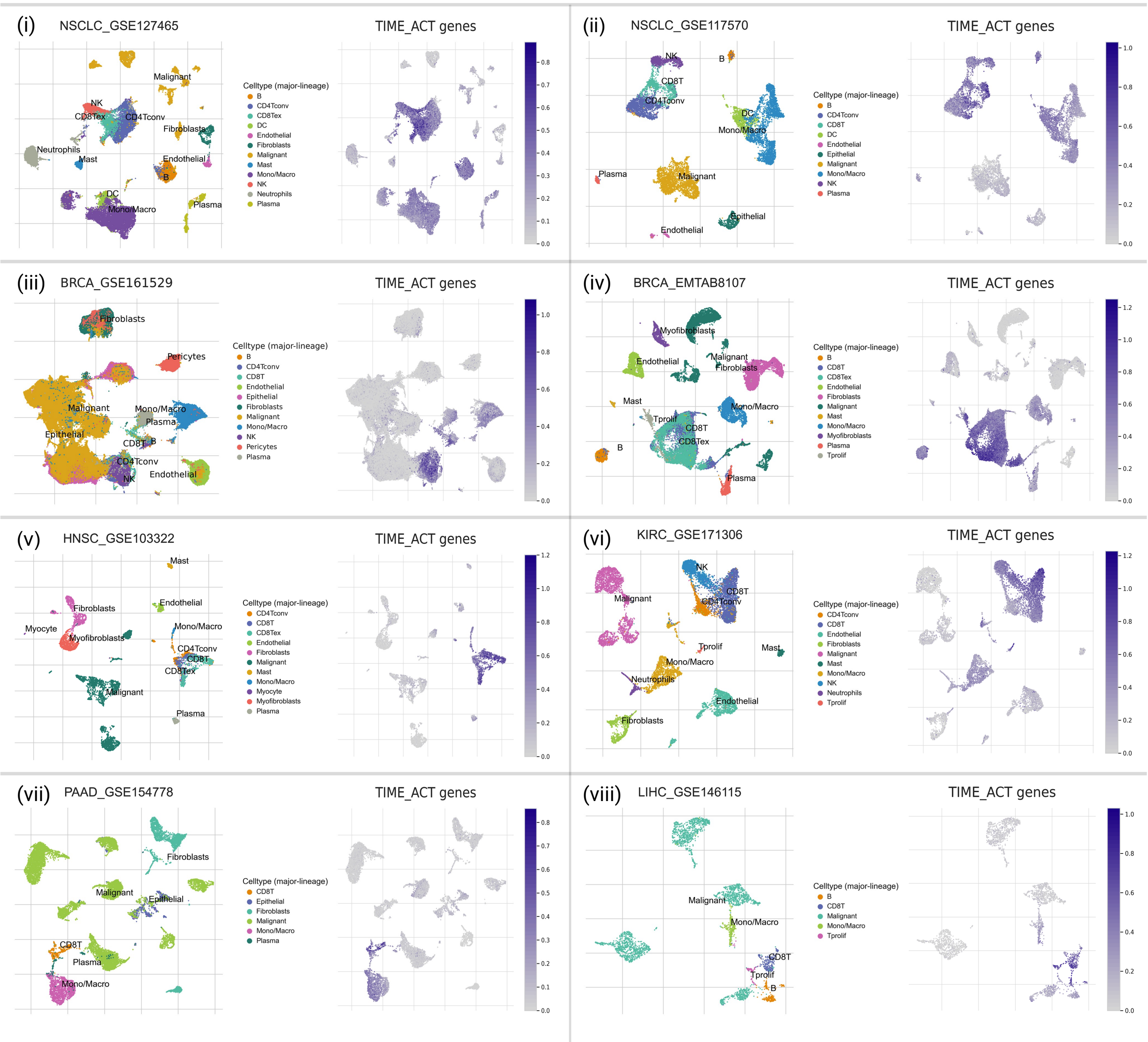
